# Anatomical registration of intracranial electrodes. Robust model-based localization and deformable smooth brain-shift compensation methods

**DOI:** 10.1101/2023.05.08.539503

**Authors:** Alejandro Omar Blenkmann, Sabine Liliana Leske, Anaïs Llorens, Jack J. Lin, Edward Chang, Peter Brunner, Gerwin Schalk, Jugoslav Ivanovic, Pål Gunnar Larsson, Robert Thomas Knight, Tor Endestad, Anne-Kristin Solbakk

**Affiliations:** Department of Psychology, University of Oslo, Norway; Department of Musicology, University of Oslo, Norway; RITMO Centre for Interdisciplinary Studies in Rhythm, Time, and Motion, University of Oslo, Norway; Department of Neurology and Center for Mind and Brain, University of California, Davis, USA; Department of Neurological Surgery, University of California, San Francisco, USA; Department of Neurology, Albany Medical College, Albany, NY, USA; National Center for Adaptive Neurotechnologies, Albany, NY, USA; Department of Neurosurgery, Oslo University Hospital, Norway; Department of Psychology and the Helen Wills Neuroscience Institute, University of California, Berkeley, USA; Department of Neuropsychology, Helgeland Hospital, Mosjøen, Norway; Tianqiao and Chrissy Chen Institute, Chen Frontier Lab for Applied Neurotechnology, Shanghai, China; Fudan University/Huashan Hospital, Department of Neurosurgery, Shanghai, China

**Keywords:** stereo electroencephalography (SEEG), electrocorticography (ECoG), intracranial EEG (iEEG), depth electrodes, subcortical grids, subdural grids, simulations.

## Abstract

Precise electrode localization is important for maximizing the utility of intracranial EEG data. Electrodes are typically localized from post-implantation CT artifacts, but algorithms can fail due to low signal-to-noise ratio, unrelated artifacts, or high-density electrode arrays. Minimizing these errors usually requires time-consuming visual localization and can still result in inaccurate localizations. In addition, surgical implantation of grids and strips typically introduces non-linear brain deformations, which result in anatomical registration errors when post-implantation CT images are fused with the pre-implantation MRI images. Several projection methods are currently available, but they either fail to produce smooth solutions or do not account for brain deformations.

To address these shortcomings, we propose two novel algorithms for the anatomical registration of intracranial electrodes that are almost fully automatic and provide highly accurate results. We first present *GridFit,* an algorithm that simultaneously localizes all contacts in grids, strips, or depth arrays by fitting flexible models to the electrodes’ CT artifacts. We observed localization errors of less than one millimeter (below 8% relative to the inter-electrode distance) and robust performance under the presence of noise, unrelated artifacts, and high-density implants when we ran ∼6000 simulated scenarios. Furthermore, we validated the method with real data from 20 intracranial patients.

As a second registration step, we introduce *CEPA,* a brain-shift compensation algorithm that combines orthogonal-based projections, spring-mesh models, and spatial regularization constraints. When tested with real data from 15 patients, anatomical registration errors were smaller than those obtained for well-established alternatives. Additionally, *CEPA* accounted simultaneously for simple mechanical deformation principles, which is not possible with other available methods. Inter-electrode distances of projected coordinates smoothly changed across neighbor electrodes, while changes in inter-electrode distances linearly increased with projection distance. Moreover, in an additional validation procedure, we found that modeling resting-state high-frequency activity (75-145 Hz) in five patients further supported our new algorithm.

Together, *GridFit* and *CEPA* constitute a versatile set of tools for the registration of subdural grid, strip, and depth electrode coordinates that provide highly accurate results even in the most challenging implantation scenarios. The methods presented here are implemented in the iElectrodes open-source toolbox, making their use simple, accessible, and straightforward to integrate with other popular toolboxes used for analyzing electrophysiological data.

## 1. INTRODUCTION

Most human intracranial electroencephalography (iEEG) studies are conducted in patients with drug-resistant epilepsy where subdural grids and depth electrodes are implanted to identify and characterize the brain areas involved in the onset and propagation of seizures (Parvizi & Kastner, 2018). iEEG records the brain’s electrical activity with a higher simultaneous spatial and temporal resolution than non-invasive methods such as scalp EEG, functional magnetic resonance imaging (fMRI), or near-infrared spectroscopy.

One critical precondition for this superior spatial resolution is the precise localization of the intracranial electrodes with respect to the brain’s anatomy. In this way, the precision and accuracy of intracranial electrode localization procedures have a crucial impact on the interpretation of iEEG in the context of clinical and cognitive neuroscience studies.

Specific patterns of neural activity in the time and frequency domains are used to define the epileptogenic zone, and the brain areas to be resected (Lesser et al., 2010; Grinenko et al., 2018; Campora et al., 2019; Dellavale et al., 2020). Additionally, motor, sensory, and language areas are usually mapped for clinical reasons, i.e., the identification and preservation of eloquent cortex (Jayakar et al., 2016; Parvizi & Kastner, 2018). Moreover, iEEG recordings provide a unique opportunity to study normal brain function during resting state and cognitive tasks with direct access to the cortex and deep structures (Jacobs & Kahana, 2010; Mukamel & Fried, 2012). iEEG signals reflect the complex interactions of diverse and distributed populations of neurons in the brain. In addition, the method allows the investigation of information flow in neurocognitive processes by modeling communication across brain areas or induced electrical stimulation activity (Collavini et al., 2021; Johnson et al., 202, Parvizi & Kastner, 2018).

Remarkably, iEEG can reliably capture high-frequency activity (HFA, 70-200 Hz), which corresponds to the average spiking activity of neurons and dendritic potentials within ∼5 millimeters of the recording electrode (Ray & Maunsell, 2011; McCarty, 2021, Leszczynski et al., 2020). Thus, high-frequency activity is an accurate marker of task-related cortical activity with high spatial accuracy (Helfrich & Knight, 2016; Holdgraf et al., 2016; Zheng et al., 2017; Parvizi & Kastner, 2018; Blenkmann et al., 2019; Johnson et al., 2020; Hamilton et al., 2021).

Over the past decade, several approaches have been proposed to register the anatomical coordinates of intracranial electrodes. Most of them are based on CT and MRI images, with a few approaches focusing on post-implantation CT and MRI images (Blenkmann et al., 2015; LaViolette et al., 2011; Hinds et al., 2018). However, the vast majority of methods use post-implantation CT and pre-implantation MRI (Hermes et al., 2010; Dykstra et al., 2012; Brang et al., 2016; Princich et al., 2013; Branco et al., 2018a; Centraccio et al., 2021; Hamilton et al., 2017, Trotta et al., 2018). In these approaches, electrodes are localized within CT images and then transferred to the co-registered MRI for anatomical description. The localization procedure is usually manual, where the coordinates are obtained by visually detecting high-intensity CT artifacts produced by the metallic contacts within a lower-intensity background (Hermes et al., 2010; Dykstra et al., 2012; Stolk et al., 2018). More recently, the localization has been approached by semi-automatic techniques such as clustering voxels of high-intensity value (Blenkmann et al., 2017; Brang et al., 2016; Branco et al., 2018a; Taimouri et al., 2013; Qin et al., 2017), interpolating coordinates given entry and target points in depth electrodes (Li et al., 2020; Arnulfo et al., 2015; Narizzano et al., 2017), using shape analysis (Centracchio et al., 2021), or deep learning (Vlasov et al., 2022).

Alternative localization approaches have been proposed based on intraoperative photography and X-ray projections (Dalal et al., 2008), intraoperative photography only (Pieters et al., 2013), solely MRI images (Yang et al., 2012), clinical neuronavigational data (Gupta et al., 2014), or electrophysiological data (Branco et al., 2018b).

Over the last few years, the spatial resolution of grids and depth electrodes has reached inter-electrode distances (IED) of 2-3 mm, and will likely improve even more (Gupta et al., 2014; Chang, 2015; Martin et al., 2018; Erhardt et al., 2020). High-density electrode arrays are more informative than low-density arrays in cognitive (Gupta et al., 2014; Jiang et al., 2018) and clinical research (Stead et al., 2010), but their localization requires enhanced spatial accuracy as well. Their reduced size presents an obstacle for the most frequently used localization algorithms because artifacts of neighboring electrode contacts cannot be disentangled from each other (Branco et al., 2018a; Hamilton et al., 2017; Narizzano et al., 2017).

Overlapping grids, clips, and other metallic objects can also challenge the detection of electrodes’ CT artifacts. Artifacts from such objects are usually treated manually (Blenkmann et al., 2017; Branco et al., 2018a; LaPlante et al., 2016; Taimouri et al., 2014) or excluded from the analysis (Brang et al., 2016).

In summary, although recent techniques have made progress in the automatic or semi-automatic localization of electrode CT artifacts, the presence of a low signal-to-noise ratio as in high-density arrays, or overlapping electrodes, cables, or artifacts still pose challenges to current methods.

In addition to the challenge of localizing for CT artifacts, other corrections might be necessary before obtaining the final anatomical coordinates. In the particular case of grid and strip electrodes, the surgical implantation procedure results in brain tissue deformation. Deformations up to 10-20 mm can occur on the brain surface around the electrodes or in deeper brain structures, precluding accurate localization directly from intracranial photographs or post-implantation CT images (Studholme et al., 2000; LaViolette et al., 2011; Brang et al., 2016). Therefore, CT localized coordinates need to be back-projected from the post-implantation space to the non-deformed pre-implantation MRI brain surface (more specifically, a smooth cortical envelope, SCE). Since brains are compressed during implantation, electrode array coordinates projected to the SCE are expected to expand (i.e., to cover more surface) to compensate for this behavior. The shape or IED of the arrays is not guaranteed to be preserved in the back transformation.

Subdural grid and strip electrode locations require compensation, but brain shifts from only stereotactic implantation of depth electrodes are less pronounced and frequent, and corrections are typically unnecessary (Elias et al., 2007). Nevertheless, less frequent cases of depth electrodes simultaneously implanted with grids or strips require corrections (Blenkmann et al., 2017; Lee et al., 2022).

The most popular brain-shift compensation methods can be grouped into two families: i) the *“Orthogonal projection”* family of methods, where electrodes are projected to the SCE surface in the direction orthogonal to the local grid surface (Hermes et al., 2010), or orthogonal to the SCE surface (Kubanek & Schalk, 2015), or in the electrodes’ principal axis (Brang et al., 2016); and ii) the *“Spring mesh”* family of methods, where the resulting coordinates are obtained while minimizing the overall deformation (Dykstra et al., 2012; Trotta et al., 2018).

Although widely used, both families come with advantages and disadvantages. In the *Orthogonal projection* family, the projections follow deformation principles that compensate for brain compressions (typically the expansion of grids when back-projecting). Still, projections are computed for each electrode individually and are susceptible to local inaccuracies, producing irregularities in their spatial distribution. On the other hand, the *Spring mesh* family of methods provides very regular and smooth results, but these methods do not compensate for the post-implantation compression of the brain. Deciding which algorithm to use in a particular case is difficult. To date, there are no clear research findings informing which method performs better for a specific brain deformation, implanted brain area, or time since implantation, to mention just a few relevant variables.

In this paper, we present two novel methods to attain high accuracy and precision in the anatomical registration of intracranial electrodes. First, we introduce a robust model-based CT artifact electrode localization technique named *GridFit*. It simultaneously localizes all the electrodes in grids, strips, or depth arrays using flexible models fitted to the CT artifacts. The algorithm has been conceptually developed to address difficult localization scenarios such as low signal-to-noise ratio, overlapping electrodes or cables, or high-density electrode arrays. We show how model parameters and performance were obtained using: i) a simulation-based approach; and ii) the application of *GridFit* to real patient iEEG data.

The second method, the Combined Electrode Projection Algorithm (*CEPA*), addresses the back-projection to pre-implantation space. It combines orthogonal projection techniques with spring mesh models and spatial regularization constraints. We compared the algorithm with well-established methods via localization error distance and modeling of electrophysiological activity. Both methods (*GridFit* and *CEPA*) are available in the iElectrodes open-source toolbox (https://sourceforge.net/projects/ielectrodes/), ensuring user-friendly access to precise anatomical registration.

## 2. MATERIALS and METHODS

We propose two novel algorithms, *GridFit* and *CEPA*. *GridFit* localizes electrode coordinates derived from CT artifacts, and *CEPA* projects the coordinates to the cortical surface of the pre-implantation MRI, compensating for brain-shift deformations. Both are integrated into the processing pipeline for intracranial electrode localization in the iElectrodes toolbox (Figure 1; Blenkmann et al., 2017). Briefly, the pipeline takes the following steps: First, presurgical MRI and post-implant CT images are co-registered. Then, to obtain the location of each electrode, CT artifacts corresponding to each array of electrodes are processed with *GridFit*. Subsequently, grid and strip electrodes are projected back to the Smoothed Cortical Envelope surface using *CEPA*. Subsequently, electrode coordinates can be visualized in 2D or 3D space and exported in formats compatible with popular analysis toolboxes, such as Fieldtrip or EEGLAB (Delorme & Makeig, 2004; Oostenveld et al., 2011). If comparisons of brain activity patterns across patients are desired, coordinates can be projected to volume-based normalized brain spaces (Ashburner & Friston, 2005) or the normalized unfolded cortical surface (Fischl et al., 1999). In the following subsections, we describe each of these steps in more detail.

**Figure 1.**
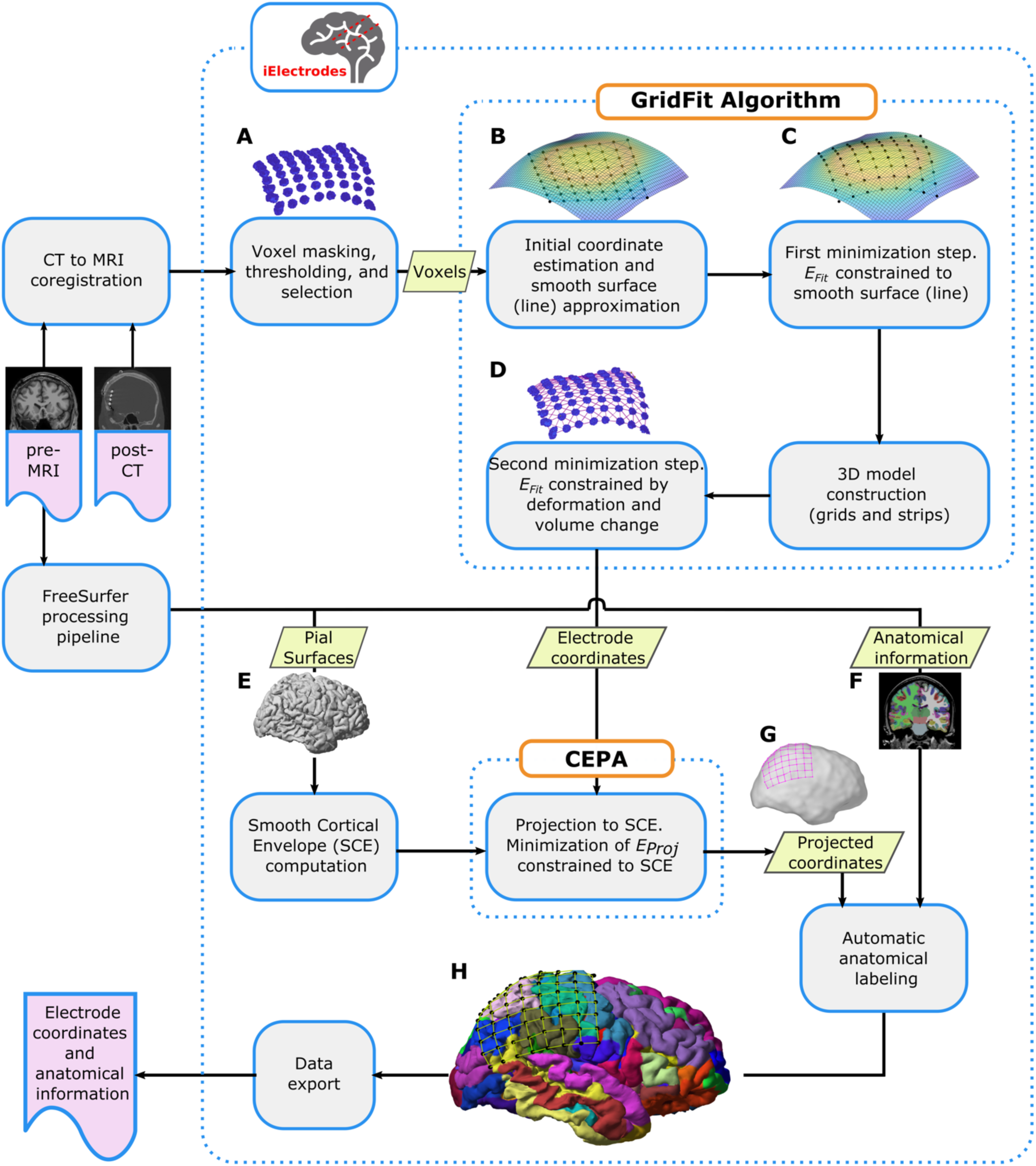
Overview of the analysis pipeline. Overview of the complete processing pipeline for localizing subdural grids, strips, and depth electrode arrays. Briefly, voxels corresponding to electrode artifacts (**A**) are extracted from the co-registered post-implantation CT image (post-CT) and processed with the GridFit algorithm (**B**, **C,** and **D**) to obtain the electrode coordinates. Pial surfaces (**E**) and anatomical information images (**F**) are derived from the pre-implantation MRI (pre-MRI) by using Freesurfer. The SCE surface is computed from the pial surface, and grid and strip electrode coordinates are projected to it using CEPA, correcting for brain-shift deformations (**G**). Depth electrode coordinates are also corrected when these arrays are simultaneously implanted with grids and strips. Finally, projected coordinates are visualized in relation to the brain anatomy incorporating the respective anatomical labels (H), and data are exported in a standardized common format. Readers are referred to the methods section for technical details about GridFit, CEPA, and other processing steps. SCE: Smooth Cortical Envelope.

### 2.1. Patients

To evaluate the performance of GridFit and CEPA on real data, we evaluated their application to iEEG data from 20 patients (10 female, mean age 33 years) with drug-resistant epilepsy in our database who were candidates for resective surgery (Kwan et al., 2010; Jayakar et al., 2016). To benchmark the two new proposed algorithms for potential problematic cases, we selected data using the following criteria: the presence of arrays with IED less than or equal to 6 mm, low SNR in the CT images, or overlapping electrodes, cables, or other artifacts, i.e., all selected patients selected meet at least one these criteria. Intracranial electrodes were temporarily implanted, and iEEG and video were continuously recorded as part of the presurgical evaluation to localize the epileptogenic focus. Patient recordings took place at four hospitals: Albany Medical College (S1-S6, S19), University of California, San Francisco (UCSF) Hospital (S7-S9), University of California, Irvine Medical Center (UCI, S10-S13), and Oslo University Hospital (OUH, S14-S18, S20).

### 2.2. Implanted electrodes

Of the 20 patients, 15 (S1-S13, S19, S20) were implanted with traditional clinical grids with 5, 6, 7, and 10 mm IED, or high-density (HD) grids with 3 and 4 mm IED (Ad-Tech Medical, USA or PMT Corporation, USA). Seven (S5, S6, S8-S10, S12, S20) were simultaneously implanted with 5 mm IED depth electrodes (Ad-Tech Medical, USA). The remaining five patients (S13-S18) were only implanted with HD, 3.5 mm IED, depth electrodes (DIXI Medical, France). Supp. Table 1 provides an overview of the implanted electrodes and their locations per patient.

### 2.3. Acquisition and preprocessing of CT and MR images

Pre-implantation MRI and post-implantation CT images were acquired as part of the clinical procedure. Patients underwent CT scans with a resolution 0.5 mm (Aquilion one, Toshiba, Japan). MRI was acquired in a 1.5 or 3 T scanner with a spatial resolution of 1 mm (Achieva MRI scanner, Philips, Eindhoven). We followed a standard procedure to localize intracranial electrodes (Blenkmann et al., 2019; Stolk et al., 2018). Pre-implantation MRI images were processed using the FreeSurfer standard pipeline (Dale, Fischl, & Sereno, 1999), obtaining individual brain segmentation images, pial surfaces (Figure 1E), curvature surfaces, and cortical parcellation images (Figure 1F). We automatically segmented the cortical surface into 74 areas per hemisphere, providing an accurate anatomical description of the cortex (Destrieux et al., 2010). Post-implantation CT images were co-registered to the pre-implantation MRI using the normalized mutual information algorithm provided by SPM (Studholme, Hill, & Hawkes, 1999). Images and surfaces were loaded into the iElectrodes toolbox and resampled to 0.5 mm resolution using fourth-order polynomial interpolation. Smooth cortical envelope (SCE) surfaces were computed for each patient’s left and right hemispheres (Figure 1G). The SCE surfaces were computed by enclosing the corresponding pial surface with a sphere of 50 mm radius using a morphological closing operation (Brang et al., 2016; Blenkmann et al., 2021).

### 2.4. Voxel masking, thresholding, and selection

To localize electrode coordinates, a cloud of CT voxels representing the grid, strip, or depth electrode array has to be selected. In this operation, a binary mask of the brain (obtained from the Freesurfer pipeline) is applied to the CT image and then eroded or dilated multiple times to remove the skull. At the same time, threshold values are manually adjusted until clusters of high-intensity CT voxels visually represent the electrodes (also known as CT artifacts; for details on adjusting threshold see Blenkmann et al., 2017). Finally, the relevant voxels of a whole array are visually selected using an incorporated drawing (brush) tool provided within the Graphical User Interface (GUI) of the iElectrodes toolbox (Figure 1A). This procedure has proved robust in localizing electrodes in previous studies (Blenkmann et al., 2017).

It is important to note that coarse selections of all the voxels from an array are taken, i.e., not a contact-by-contact selection, providing a fast and replicable process. These selections might include artifacts from cables, contacts corresponding to overlapping arrays, or blurred, not-well-defined, artifacts from the contacts of interest. In low SNR conditions, electrodes cannot be isolated when adjusting the threshold and might stay connected with their neighbors.

This procedure results in a set of *N_Vox_* thresholded voxels representing an electrode array. It consists of voxel coordinates ***v****_n_* and voxel intensities *w_n_,* where *n = 1, …, N_Vox_*. The intensity values within the array are normalized between zero and one. This set of voxel coordinates and intensities serves as the *GridFit* algorithm’s input data to localize the electrode coordinates ***e*** of an *N_Rows_ x N_Cols_* array (*N_Elec_* electrodes in total) and IED *D*. As an optional step, after the voxel selection procedure, a set of “fixed” electrode coordinates ***e_Fix_*** can be visually selected at the center of the CT artifacts. These coordinates can be used if the *GridFit* algorithm does not converge to a feasible solution.

### 2.5 2D and 3D electrode array models

Model-based localization (*GridFit*) and the brain-shift compensation algorithm (*CEPA*) require 2D and 3D models of the arrays. 3D models are exclusively used in the second step of *GridFit* for grids and strips. Therefore, we present the 2D and 3D models before the *GridFit* and *CEPA* methods.

The number of rows and columns (*N_Rows_ x N_Cols_*), the IED *D*, and the thickness *T* (3D models only) are used to define their geometrical characteristics. Models are built by defining points and their connections (Figure 2A). Auxiliary structural model points ***m*** are defined to provide additional spatial structure. Some of these points are located at the electrode coordinates ***e***; therefore, the *N_Elec_* electrode coordinates are a subset of the *N_Mod_* structural model points (please see below how subsets are defined). We describe ***e*** and ***m*** points separately to simplify the mathematical notation. However, since ***e*** is a subset of ***m***, modifying any of the points in common alters both sets simultaneously.

**Figure 2.**
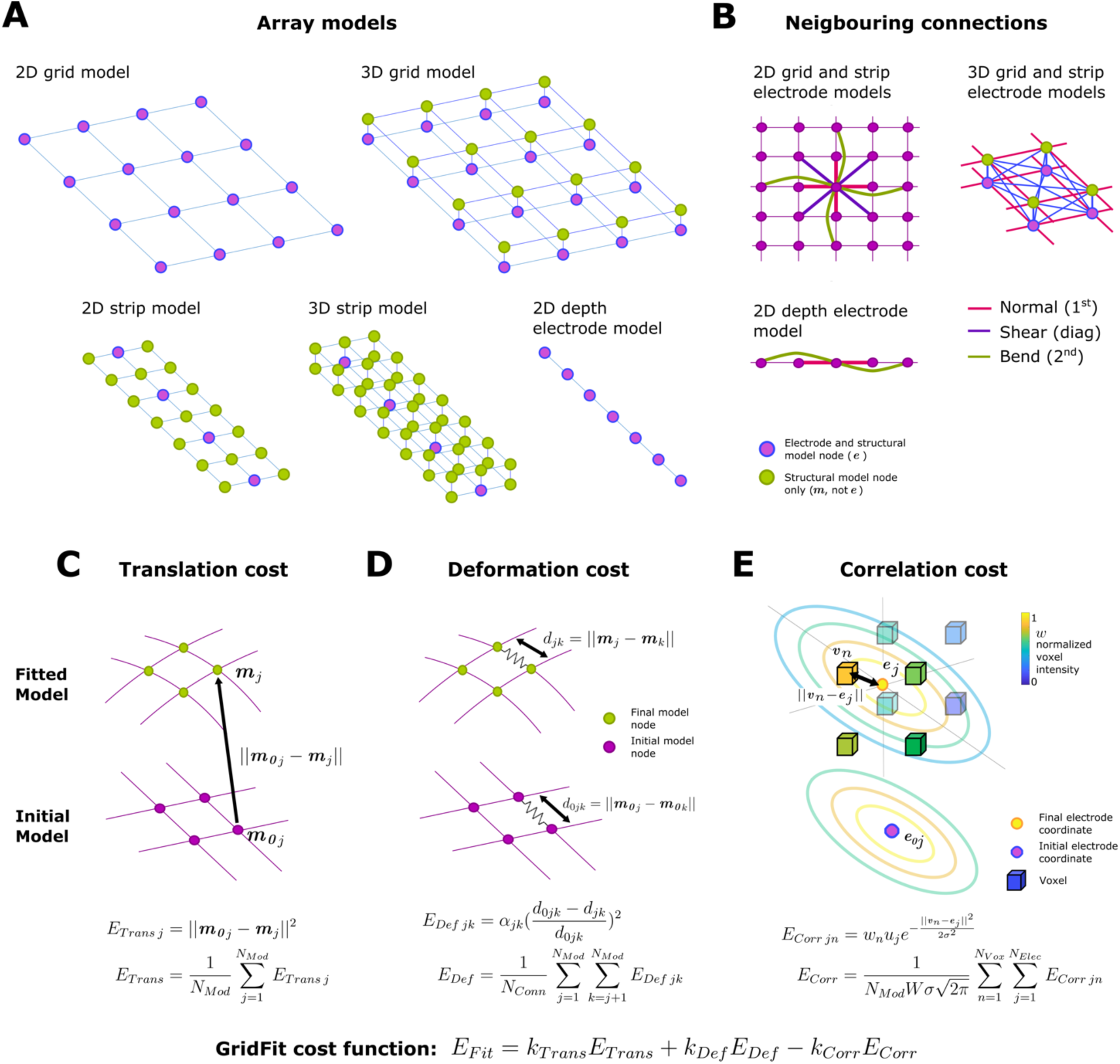
Array models and GridFit cost function sub-components. **A.** Variety of 2D and 3D models used in the GridFit algorithm. Note that the electrode coordinates (magenta) are a subset of the model node coordinates (green). **B.** Diagram showing the neighboring connections for the 2D and 3D models. **C, D,** and **E.** Diagrams representing the different components of the cost function minimized in the GridFit algorithm. The final model coordinates (green) are depicted on top of the initial model coordinates (magenta). For illustrative purposes, only 2D models are shown. **C.** The translation of model points from the initial to the final location increases ETrans proportionally to the square of the distance. **D.** The model deformation, either by contraction, expansion, shearing, or bending, affects the distance between model nodes and increases the deformation cost EDef. **E.** An electrode coordinate in the vicinity of voxels corresponding to an electrode artifact increases the correlation cost ECorr. Color-coded ellipsoids depict how the Gaussian function decreases with the electrode coordinates’ distance. For illustrative purposes, only first neighbor connections are shown in **A**, **C**, **D**, and **E**. For a description of the formulas, we refer the reader to subsection 2.6.3.

Connections are defined between neighboring structural points. Neighbors are defined in three ways: first neighbors, if orthogonally adjacent; second neighbors, if separated by one point orthogonally; and diagonal neighbors, if diagonally adjacent (Figure 2B). The 2D and 3D models used for the *GridFit* and *CEPA* algorithm are defined as follows:

● 2D models are built for

○ Grids: by connecting each point ***m****_j_* with its first, second, and diagonal neighbors, where electrode coordinates and structural points are the same (*N_Elec_ = N_Mod_*, ***e*** *= **m)***.
○ Strips: by extending the array’s narrowest dimension by half of the IED on each side. A lattice grid of points ***m*** is constructed at D/2 resolution. The first, second, and diagonal neighbor model points are connected. Electrode locations ***e*** correspond to the central subset of coordinates from ***m***.
○ Depth electrodes: by connecting the first and second neighbor points, and consisting of structural points ***m*** at each electrode location ***e (****N_Elec_ = N_Mod_, **e** = **m)***.
● 3D models for grids and strips are built by positioning two 2D structural grid models on top of each other, separated by the array thickness. Each model point ***m*** is connected to all 1^st^ and diagonal neighbors in the 3D grid (no second neighbor connections). The electrode coordinates ***e*** are assigned to only the subset of ***m*** coordinates on the bottom layer (i.e., ***e*** *= **m*** for the bottom layer), while the top is used for structural reasons (i.e., only ***m*** points).

### 2.6. *GridFit* algorithm: Automatic localization of CT artifacts

*GridFit* is a model-based algorithm that uses the information from all voxels to reconstruct the coordinates of an electrode array (either depth, grid, or strip). Briefly, the approach is implemented by fitting a flexible array model to the set of selected CT artifacts (*N_Vox_* voxels at ***v*** with intensities ***w***) in a two-step cost-minimization approach. The central idea of the method is to penalize model deformations during the fitting processes while rewarding the proximity of candidate coordinates to clusters of voxels representing electrodes.

The first cost-minimization step is constrained to a surface or line describing the main location of voxels and provides a coarse approximation to the final electrode location. The second cost-minimization step is not constrained to any surface and uses more sophisticated models, providing better control of the final localization. The diagram in Figure 1 shows the main steps of the algorithm, which are described in more detail in the following subsections.

#### 2.6.1. PCA rotation and smooth surface (line) approximation of the CT artifacts

To simplify coordinate handling in the subsequent steps, we first rotated the coordinate system of the selected CT voxels using principal component analysis (PCA). The new x-, y-, and z-axes represent PC1, PC2, and PC3, respectively, accounting for high to low data variance. At the end of the *GridFit* algorithm, the localized coordinates are rotated back to the original space.

The procedures described below require an approximation function of the voxels, which is computed as follows. For grids, a surface function *z = f_surf_(x, y)* is constructed by fitting a smooth surface to the set of voxel coordinates ***v*** (Figures 1B & 1C; Locally Weighted Scatterplot Smoothing algorithm; Cleveland, 1979).

For depth electrodes and strips, instead, a unidimensional function *y = f_lin_(x)* is built by fitting the first two dimensions of the voxel coordinates ***v*** with a 3rd-order polynomial function. Due to its small variance, the last dimension is ignored.

#### 2.6.2. Initial coordinate estimation

An initial set of electrode coordinates ***e_0_*** (e.g., Figure 1B) is needed as an input for the first cost-minimization procedure. These coordinates are obtained following a procedure described in Blenkmann et al. (2017). For grid arrays, voxels ***v*** are projected onto the first two components of the PCA space defined in the previous section. Subsequently, a convex hull is iteratively trimmed until the four corners of the grid are isolated. These corners are then back-projected to the 3D space using *f_surf_*. Finally, the non-corner coordinates are linearly interpolated. For depth electrode arrays, the initial coordinates ***e_0_*** are estimated by uniformly distributing the electrodes between the extremes of the first PCA component of the voxel coordinates ***v*** and back-projecting them to the 3D space using *f_lin_*.

#### 2.6.3. GridFit cost function

*GridFit* was conceived as a model-based approach to localize CT artifacts in cases where the information is noisy or missing. In this way, the algorithm does not localize individual electrodes but simultaneously localizes all *N_Elec_* electrodes within an array. We formulated the localization as an optimization problem to achieve our objective, i.e., finding an optimal set of coordinates ***m*** that minimizes a cost function. In our algorithm, the optimal solution minimizes the displacement and deformation of the array while maximizing the correlation between electrode locations (***e***) and thresholded voxels (***v***, ***w***). *GridFit* uses a two-step optimization, i.e., the same cost function is minimized two times under different sets of constraints for each step (see details in sections 2.6.4 and 2.6.5). The first minimization gives a coarse and fast set of coordinates that serve as starting points for the second minimization, where more precise coordinates are obtained. The respective cost function to be minimized is

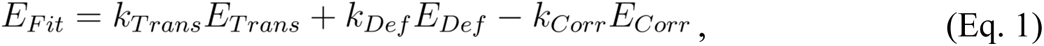

where *E_Trans_*, *E_Def,_* and *E_Corr_* are the translation, deformation, and co-registration cost functions, accompanied by *k_Trans_*, *k_Def,_* and *k_Corr_*, the translation, deformation, and co-registration constants, respectively (the procedure to obtain optimal parameter values is discussed in section 2.7).

The translation cost function *E_Trans_* is defined as

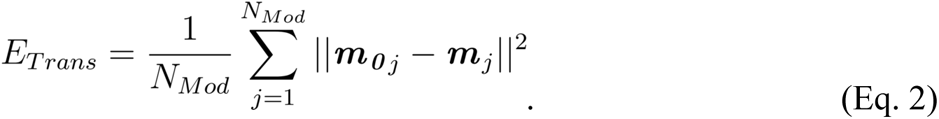

This function penalizes the translation of the structural points to the actual location ***m*** from their initial location ***m_0_***, and therefore an appropriate initial location ***m_0_*** (including ***e_0_***) is needed (Figure 2C).

*E_Def_*, the deformation cost function, is defined as

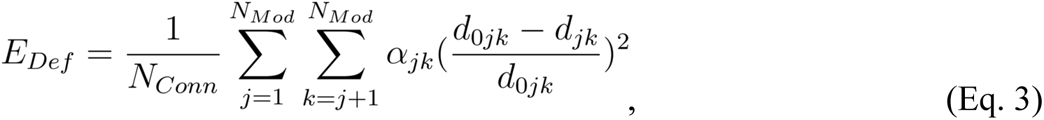

where *N_Conn_* is the total number of neighbor connections in the model, *α_jk_* is a parameter which equals 0 for non-neighboring points and a weight value (in the 0-1 range) for neighboring points, *d_jk_* is the Euclidean distance between structural points *j* and *k* defined as *d_jk_* = ||*m_j_* − *m_k_*||, and *d_0 jk_* is the distance between points *j* and *k* in the 2D or 3D initial model *d*_0*jk*_ = ||*m_0j_* − *m_0k_*||. The deformation cost function *E_Def_* considers the total network deformation where the network is built with the equivalent of “springs” connecting neighboring points (Figure 2D). When the grid is deformed (compressed, stretched, sheared, or bent), the deformation cost added is proportional to the square difference between the initial and actual distances in each spring connecting pairs of structural points.

Finally, the co-registration cost function *E_Corr_* considers the complete array of *N_Elec_* electrodes in relationship with the *N_Vox_* thresholded voxels and increases its value as their location matches. Note that the minus sign preceding the co-registration term in Eq. 1 imposes that the optimization algorithm will attempt to maximize *E_Corr_*. The co-registration cost function *E_Corr_* is defined as

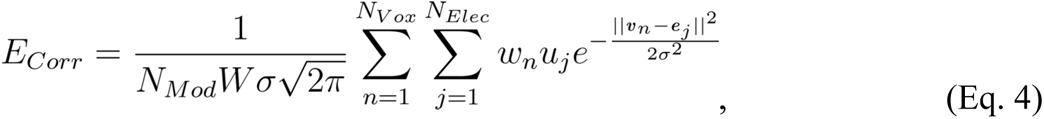

where *σ* is a regularization parameter of the spatial dispersion,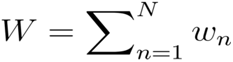, and *u_j_* is the ratio of connections associated with electrode *j*. The co-registration cost function *E_Corr_* is composed of Gaussian functions that depend on the distance between voxels and electrode*s*, i.e., *||**v**_n_ - **e**_j_||*, weighted by the voxel intensity *w_n_*. Voxels with a high-intensity value will contribute more, as they may contain more reliable information about the electrode locations than those with lower values. *σ* defines the spatial dispersion (standard deviation) of the Gaussian function, i.e., the spatial sharpness with which voxels around an electrode are considered. Consequently, when an electrode location reaches a cluster of voxels representing an electrode, the value of the function *E_Corr_* increases (Figure 2E). The variable *u_j_* reduces the co-registration effect on the edge electrodes and increases the effect on the inner electrodes. This correction is to counterbalance the *E_Def_*’s larger effect on inner electrodes compared to bordering ones, given their presence in more connections.

Altogether, the minimization of the *E_Fit_* function will fit an array of electrodes to a selection of voxels, minimizing the displacement and deformation of the array, and maximizing the correlation between electrode locations and thresholded voxels.

#### 2.6.4. First minimization step

In the first cost-minimization procedure, 2D models are used to provide a rough estimate of the electrode locations. The cost function *E_Fit_* (Eq.1) is minimized, while the electrode locations are constrained to be located on the smooth approximation function of the CT artifacts (*f_surf_* or *f_lin_*, Figure 1C). For grids and strips, the minimization is constrained to

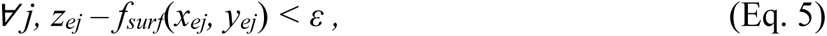

where ***e****_j_ =* [*x_ej_, y_ej_, z_ej_*] is the location of electrode *j*, and *ε* is an arbitrarily small distance. For depth electrodes, it is constrained to

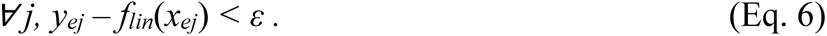

Matlab’s Interior Point algorithm (implemented in Matlab’s “fmincon” function, Waltz et al., 2005) is used to solve the cost-minimization problem, where ***e_0_*** coordinates are used as initial conditions, and a set of coordinates ***e****_First Fit_* is obtained. *σ* = *D* and *ε* = 1 mm were used.

Optimizations were terminated when the change in *E_Fit_* (Eq. 1) was less than *D* 1E-2.

#### 2.6.5. Second minimization step

The second cost-minimization procedure results in the final estimation of the electrode coordinates. ***e****_First Fit_* coordinates, the result of the first fitting step, are used as initial conditions.

3D models are created from the previously used 2D models for grids and strips. The initial coordinates ***m_0_*** are extrapolated from the 2D solutions ***e****_First Fit_.* A new 2D model layer is created on top of the previous one, at a distance *T* (thickness) between the two. Then, these 2D layers are connected (Figure 2A). 3D models were adopted for grids and strips because they offer better control of the deformations.

2D models are used again for depth electrode arrays since deformations are not pronounced and increasing the model complexity would unnecessarily raise computational demands.

The final electrode coordinates are localized (Figure 1D) by minimizing the cost function *E_Fit_* (Eq. 1), constrained to:

1. A maximum deformation between structural points, implemented as

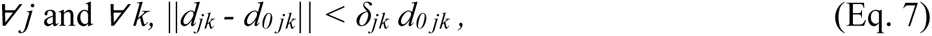

where *j = 1, …, N_Mod_, k = 1, …, N_Mod_, d_jk_* is the final distance between ***m****_j_* and ***m****_k_*, *d_o jk_* is the initial distance between the same pair of structural points. *δ_jk_* is defined for each set of points, being more strict for neighboring points (*δ_jk_ =* 10% deformation) and more relaxed for non-neighboring (*δ_jk_ =* 25% deformation), in this way providing a local and global control on the maximum deformation allowed.

2. A volume change (for 3D models only), defined as

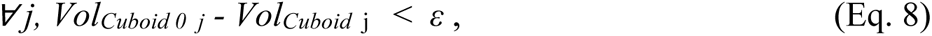

where *j=1, …, N_Cuboid_, Vol_Cuboid j_* is the volume of a rectangular cuboid *j, Vol_Cuboid 0 j_* is the initial volume of the same cuboid, and *ε* is an arbitrary small number. *N_Cuboid_* cuboids cover the complete 3D model and are built connecting the first neighbor nodes (vertices). This constraint guarantees that the resulting grid or strip volume will be the same as the initial one after the optimization.

1. A set of fixed coordinates (optional constraint), defined as

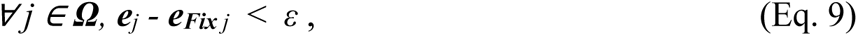

where ***Ω*** is the set of electrodes associated with a fixed coordinate (see definition in subsection 2.4). It is up to the user’s needs whether this last constraint is implemented or not.

In the following, we will refer to ***e****_GridFit_* as the set of electrode coordinates that minimizes *E_Fit_* in the second cost-minimization step. The Interior Point algorithm is used for this purpose (Waltz et al., 2005). *σ* = *D*/4 and *ε* = 0.01 mm parameters were used. Optimizations were terminated when the change in *E_Fit_* (Eq. 1) was less than *D* 1E-6.

### 2.7. Definition of GridFit parameters and localization performance evaluation using simulations

The *GridFit* algorithm encompasses several parameters that influence its behavior. Since the optimal set of parameters was unknown, we determined them using synthetic data from an open-source simulation platform (Blenkmann et al., 2022). We used more than 850 depth and 3300 grid and strip implantation scenarios. Scenarios were available with different implantation coordinates and brain curvatures, and several geometric configurations were simulated, including IEDs of 3, 5, and 10 mm, and array sizes ranging from 1 x 4 to 8 x 16 electrodes (grids and strips) or 6 to 15 aligned contacts (depth arrays).

CT artifacts were carefully modeled over the cortical surface, aligned parallel to the cortical surface (grids and strips), or within the brain parenchyma, following their principal array axes (depth arrays). For each scenario, multiple simulations were performed at several noise levels.

We simulated three levels of CT noise by spatially displacing voxels from their original positions in a random direction with Low, Medium, and High intensity (*σ* = 0.1, 0.2, and 0.4, respectively). Moreover, we also simulated overlapping grids or strips, and curved depth electrode arrays. To quantify localization accuracy, we computed the normalized median error for each array as

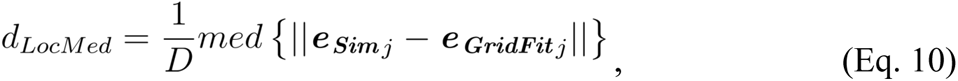

and the normalized maximum error as

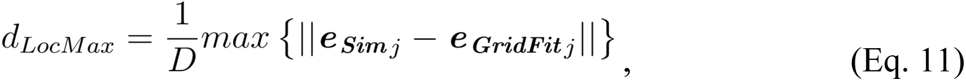

across the set of all electrodes, i.e., *j=1,.. N_Elec_*, where ***e_Sim_*** *_j_* and ***e_GridFit_*** *_j_* are the simulated and localized coordinates. We excluded simulations with *d_Loc Med_* values larger than five standard deviations from the median. Finally, we defined accuracy as *1-median(d_Loc Med_)* across arrays.

We performed three different rounds where simulated coordinates were localized. In the first round, a wide range of parameter values were evaluated within a small set of test scenarios to assess roughly their effect on the localization. In this way, we heuristically defined all but one of the parameters (*K_Corr_*), ensuring good localization results in all simulated test scenarios.

In the second round, a more detailed mapping of the effect of *K_Corr_* was obtained. To characterize the performance, between 20 and 30 simulation test scenarios were randomly performed in every combination of electrode size, curvature, IED, noise level, and overlap, making a total of ∼69000 simulations. In this round, all the constraints described in the second fitting step (subsection 2.6.6, Eq. 7, 8, and 9) were removed to reduce computational time. By measuring the median localization error, these simulated scenarios allowed us to define the optimal value of *K_Corr_* given the array geometry and the presence or absence of overlaps. Moreover, optimal *K_Corr_* values for other geometries than the simulated ones (e.g., the real cases) are obtained by interpolation, making the algorithm application range more general.

Finally, we evaluated the algorithm’s performance in a third round using optimal parameters in 16 to 60 newly and randomly generated simulations per implantation scenario (6004 simulations in total). Fewer simulations were performed on larger arrays due to computational constraints. N-way ANOVA was used to define the effect of IED, noise, overlap, array type, and interactions on the normalized median error, and ω^2^ was used to estimate effect sizes (Lakens, 2013).

The localization of simulated arrays was evaluated using near-optimal parameters (sub-optimal *K_Corr_* by one order of magnitude above and below optimal) to assess the robustness of *GridFit* to deliberate changes in the parameters. Approximately 12000 simulations were used for this purpose.

### 2.8 Validation of GridFit localization using real data

To further assess the validity of our method, we evaluated the *GridFit* localization algorithm with real imaging data. To do this, we used the optimal set of parameters obtained from the previous simulations to perform the localization in real data obtained from 20 patients.

First, *GridFit* results were evaluated by visual inspection. Localized coordinates were visually compared with the CT artifacts to detect any convergence error (outliers). Coordinates were expected to lie within the CT artifacts (when available) and to be regularly distributed covering the array. If a single electrode was mislocalized, i.e., the localized coordinate is outside the visible CT artifact, we defined the array as not successfully localized.

Then, *GridFit* localized coordinates (***e****_GridFit_*) were contrasted with a classical manual visual localization procedure by two trained users. More precisely, for the manual localization, coordinates ***e_Vis_*** were visually defined as the center of the clusters of high-intensity voxels after thresholding and masking the CT artifacts (as defined in section 2.4).

The distances between ***e****_GridFit_* and ***e_Vis_*** were used to validate our procedure. Additionally, we compared the mean absolute deviation (MAD) of the inter-electrode distances for ***e****_GridFit_* and ***e_Vis_*** to assess unexpected deformations in the geometry of localized arrays.

### 2.9. Combined Electrode Projection Algorithm (*CEPA*): Brain-shift compensation for grid and strip electrode coordinates

To compensate for the brain-shift deformation, the initial set of coordinates (i.e., the localized CT artifacts) needs to be back-projected to the SCE (Figure 1G). We intended that projections expand with the projection distance and simultaneously have spatially smooth deformations. Therefore, we approached this problem by combining orthogonal projections, spring mesh models, and regularizations accounting for the roughness of spatial deformations. The projection was formulated as aminimization problem using the previously defined 2D models (section 2.5). The function to be minimized is

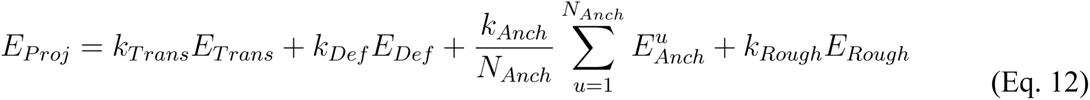

constrained to

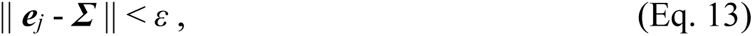

for *j=1, …, N_Elec_*, where *ε* is an arbitrary small number, and ***Σ*** is the SCE surface described as a set of vertices and faces. Figure 3 shows a schematic representation of the method.

**Figure 3.**
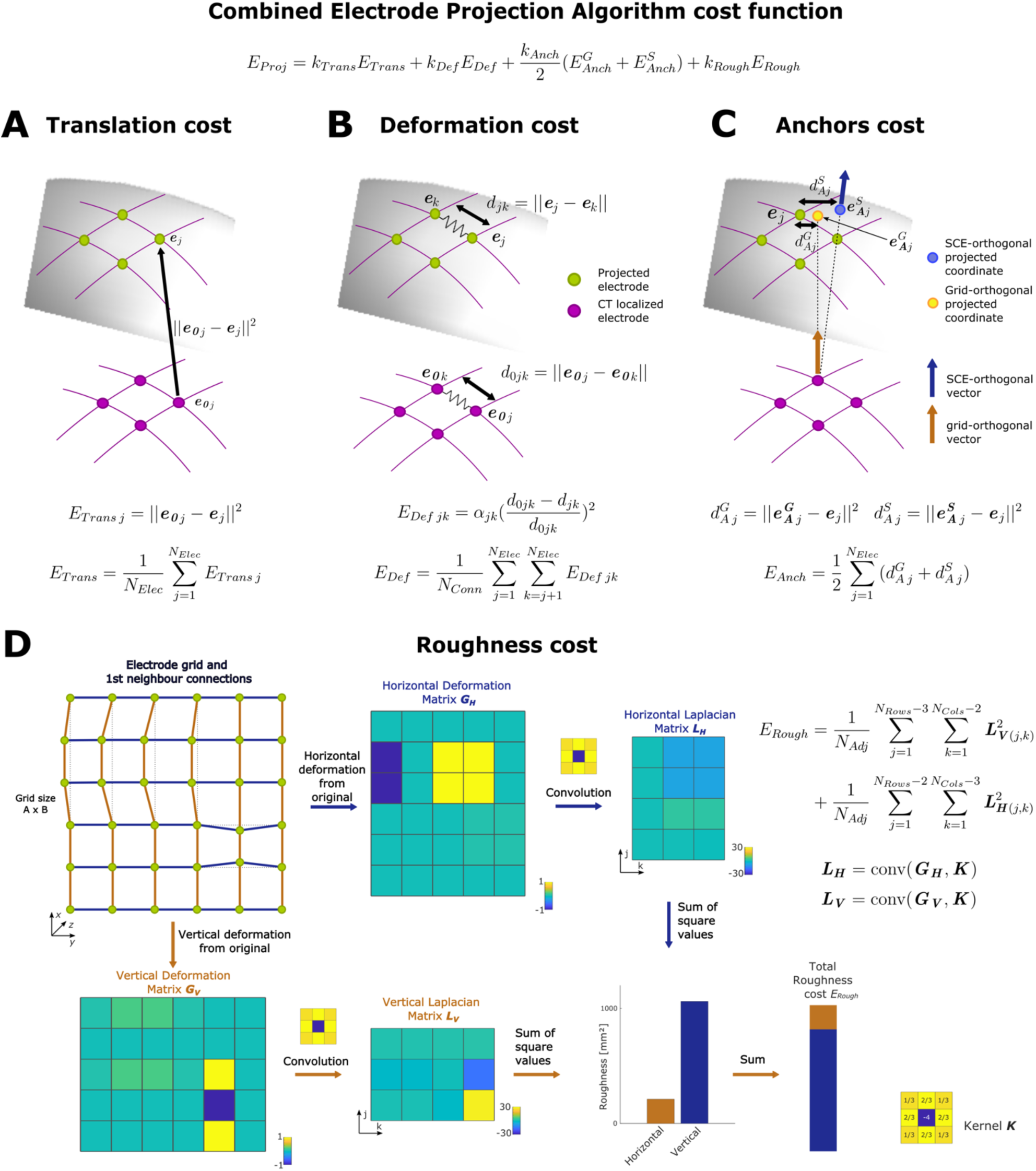
Combined Electrode Projection Algorithm (CEPA) diagram. **A.** The translation of electrode coordinates from the initial to the final location onto the SCE surface (gray surface) introduces an increase of ETrans proportional to the square of the distance. **B.** The model’s deformation, either by contraction, expansion, shearing, or bending, affects the distance between electrode nodes and increases the deformation cost EDef. **C.** The anchor’s cost EAnch increases proportionally to the distance between the anchor coordinates and the final location of electrodes. Two anchor coordinates are considered, **e_A_^S^** based on the SCE-orthogonal direction vector and **e_A_^G^** based on the grid-orthogonal direction vector. **E.** Deformations along the vertical or horizontal connections increase the roughness cost ERough proportionally to the local consistency of these deformations. In the example, a larger but homogeneous deformation in the horizontal direction introduces a relatively small roughness cost change compared with a small but highly local deformation in the vertical direction. For a complete description of the formulas, we refer the reader to section 2.9.

The first two terms in Eq. 12 are computed as those used in the *GridFit* algorithm. The cost function *E_Trans_* (Eq. 2) accounts for the translation of the coordinates from the initial to the final location (Figure 3A). The cost function *E_D_* (Eq. 3) considers the deformation introduced to a spring mesh connecting the electrodes and reflects expansions between the initial and final states (Figure 3B).

The third term of Eq. 12 describes the cost associated with deviations from anchor points. Anchor points ***e_A_****^u^* are, in general terms, defined as a set of electrode locations over the SCE surface obtained by other methods, where *u = 1, …, N_Anch_*, and *N_Anch_* is the number of anchor coordinate sets (Trotta et al., 2018). The cost function *E^u^_Anch_* associated with each anchor set *u* is defined as

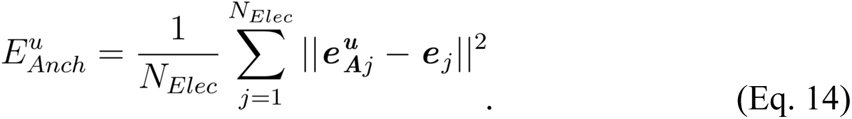

In the current *CEPA* implementation, we use two sets of anchor coordinates defined by orthogonal projection methods (*N_Anch_ = 2*). Both methods project individual electrodes onto the SCE surface given a direction vector (Figure 3C). In the first method, direction vectors are computed as the orthogonal vectors to the plane made by each electrode and its closest neighbors, as Hermes et al. (2010) proposed, producing anchored coordinates ***e_A_^G^***. This method is only available for grids. In the second method, orthogonal vectors are computed from the average orthogonal vectors in the SCE surface within a certain radius from the electrode, as described in Kubanek and Schalk (2015), producing anchored coordinates ***e_A_^S^***. The following sections will refer to these orthogonal projection methods as *“Normal-Grid”* and *“Normal-SCE”*, respectively.

The general definition of the anchoring cost term allows the expansion of the current method to other sources of information. These could include, for example, electrode coordinates obtained via intraoperative neuronavigation tools (Gupta et al., 2014), intraoperative photographs (Trotta et al., 2018), post-implantation MRI localized coordinates (Yang et al., 2012), or even methods to be developed.

The last term of Eq. 12 accounts for the spatial roughness of the deformations. Our rationale is that brain compressions are not established as abrupt changes of the cortical surface, but rather as spatially smooth compressions given the structural support provided by the brain tissue and by the grids or strips. Therefore, the progression from less to more compressed areas should be smooth, and the back-projection of the electrode coordinates to the uncompressed cortical surface should also follow a smooth spatial deformation pattern (Hartkens et al., 2003; Skrinjar et al., 2002).

Accordingly, we define the roughness cost function as

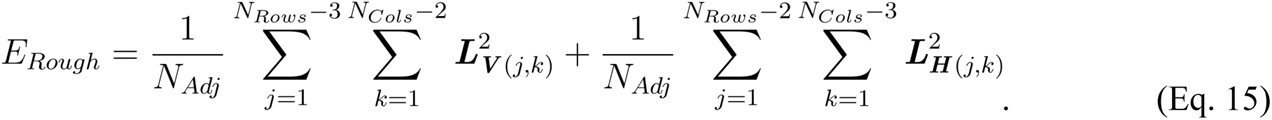

The two terms of the equation sum over the elements of 2D matrices obtained by the discrete Laplacian operator applied to the vertical and horizontal deformations. These are computed as ***L_v_*** = Δ(***G_v_***), of dimensions *N_Cols_-3 x N_Rows_-2,* and ***L_H_*** = Δ(***G_H_***), of dimensions *N_Cols_-2 x N_Rows_-3*, respectively (Figure 3D). *N_Adj_* is the number of adjacent pairs of electrodes.

For grid cases, the Laplacian operator is implemented as the 2D convolution of a nine-point stencil kernel ***K*** (3 x 3, Figure 3D, right bottom corner) with deformation matrices ***G_V_*** and ***G_H_***. The deformation matrices are obtained as the difference between the initial and final distance matrices along with the vertical or horizontal connections, i.e., ***G_V_*** = ***D_v0_*** − ***D_V_*** with dimensions *N_Cols_*-1 x *N_Rows_*, and ***G_H_*** − ***D_H0_*** − ***D_H_*** with dimensions *N_Cols_* x *N_Rows_ -1*, respectively. The 2D organization of the elements in the distance matrices follows the connection’s location within the grid along the rows and columns. For example ***D_H_***_(1,1)_ = || ***e***_(1,1)_ - ***e***_(1,2)_ || and ***D_V_****_(1,1)_* = || ***e***_(1,1)_ - ***e***_(2,1)_ ||, where ***e****_(j,k)_* denotes the coordinate of the electrode located in row *j* and column *k* of the grid. For strips and 2 or 3 -rows or -columns grid cases, we use a 1D convolution of a three-point stencil kernel (i.e., [1 -2 1]) with the deformation matrix associated with the array’s longest dimension.

Matlab’s Interior Point algorithm (Waltz et al., 2005) was used to solve the minimization problem (Eq. 12). Optimizations were terminated when the change in *E_Proj_* cost was less than 1E-6. For the *CEPA*, parameters were set to *k_Trans_* = 1, *k_Anch_ = k_Def_ = k_Rough_* = 100, and *ε* = 0.1 mm. For the *Springs* method, *k_Trans_* = 1, *k_Def_* = 1000, and *ε* = 0.1 mm (Trotta et al., 2018; Blenkmann et al., 2021).

### 2.10. Brain shift compensation of depth electrodes implanted simultaneously with subdural grids or strips

Depth electrodes simultaneously implanted with subdural grids or strips also require compensation for brain shift deformations. We approached this compensation by computing a weighted displacement field from the projected grid or strip coordinates (Taimouri et al., 2014; Blenkmann et al., 2017; Li et al., 2020). Translations are applied to the depth electrodes depending on their distance to the grid or strip electrodes. Weight functions between depth electrode *j* and grid or strip electrode *k* were defined as

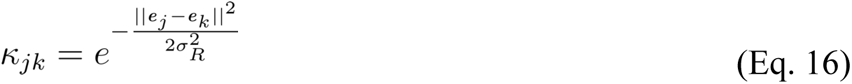

And

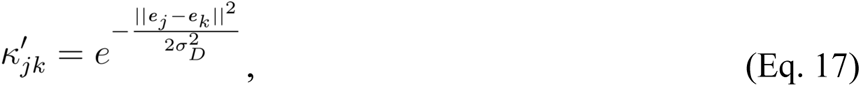

where *σ_R_* = 5 mm and *σ_D_* = 30 mm are regularization parameters (Blenkmann et al., 2017; Li et al., 2020). Finally, the translation vector applied to each implanted depth electrode *j* is computed as

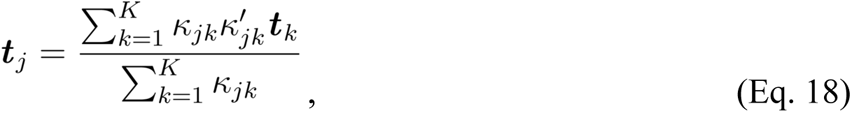

where ***t****_k_* is the translation vector applied to grid or strip electrode *k* in the back-projection procedure, i.e., ***t****_k_ = **e******_0_****_k_ - **e**_k_*, and the sum iterates over all *K* grid and strip electrodes implanted.

### 2.11. Automatic anatomical labeling

We used a probabilistic approach to assign anatomical labels to each electrode (Figure 1H). The percentage of each label is computed within a 3 mm radius volume surrounding the electrode coordinate. The anatomical description labels are obtained from FreeSurfer anatomical parcellation atlases (Desikan et al., 2006; Destrieux et al., 2010), or normalized atlases (Jenkinson et al., 2012; Yeo et al., 2011). These can be loaded by default in the iElectrodes toolbox, while others can be easily added. This approach allows labeling contacts with specific cortical denominations, subcortical structures, white matter, functional networks, or others. The labeling is relevant for describing and comparing activated areas across patients.

### 2.12. Validation of CEPA’s electrode projections on real data

We validated *CEPA* with other available electrode projection methods. First, we evaluated the error distance, roughness, and deformation of the projected coordinates by the different approaches.

Then, we compared the methods regarding how much of the local electrophysiological activity can be explained by modeling (given the projection coordinates). The next two subsections expand these ideas.

#### 2.12.1. Validation through pre-validated methods

We compared *CEPA* projected coordinates with those obtained by three previously validated methods: i) a spring mesh deformation approach introduced by Dykstra et al. (2012) and modified by Trotta et al. (2018), referred to as *“Springs”* in the following sections, and two orthogonal projection algorithms, ii) the *“Normal-Grid”* (Hermes et al., 2010), and iii) *“Normal-SCE”* (Kubanek & Schalk, 2015) methods.

Error distances were computed between the projection coordinates and reference coordinates ***e_Ref_***. By following the principle of parsimony (Occam’s razor), we defined ***e_Ref_*** as the mean of the coordinates resulting from the three pre-validated methods, assuming that they are randomly distributed around the true anatomical recording site. However, it is difficult to know to what extent the ***e_Ref_*** represents the ground truth. Consequently, we evaluated the error distances while considering each of the other three methods as a reference.

Given that projection coordinates are expected to deform smoothly (which compensates for smooth brain deformations; Hartkens et al., 2003; Skrinjar et al., 2001), we evaluated the roughness of the spatial deformation for the different back-projected coordinates using Eq. 15.

Finally, projected coordinates are expected to expand with the projection distance, i.e., cover larger areas when back-projected to compensate for stronger brain compressions during implantation. For each electrode, we defined the projection distance as the length between the localized CT artifact (***e_0_****_j_*) and the back-projected coordinate (***e****_j_*) normalized by the distance to the brain center of mass (*d_BCM_*), i.e.,

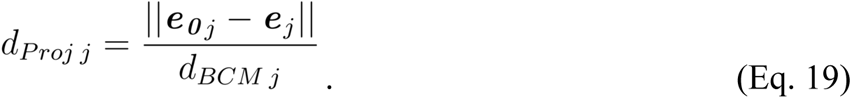

To capture the amount of expansion of grids and strips locally (at each electrode *j*), we defined the local deformation in terms of the IED *D* as

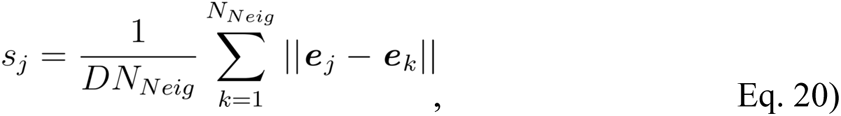

where *k* iterates over the *N_Neig_* 1^st^ neighbors around the electrode under evaluation.

In other words, local deformations (*s*) are the average inter-electrode distance change.

We evaluated how the projection distance affected local deformations in the different algorithms using Linear Mixed Effects (LME) models, with fixed effects for distance and random effects for intercept and distance grouped by electrode array and patient.

#### 2.12.2. Validation using electrophysiological data

To assess the localization accuracy independently from the CT images, we compared expected electrophysiological activity patterns given the location of the back-projected electrodes in five patients (9 grids from patients S1, S3, S10, S19, and S20; 526 channels in total). Electrodes located over gyri are closer to the cortex than electrodes located over sulci, and therefore higher intensity of neuronal activity is expected to be recorded (Bleichner et al., 2011). Accordingly, the distance of the electrodes to the cortex can partially explain the magnitude of the neural activity measured by the high-frequency activity (HFA, Branco et al., 2018 NI). In this way, we computed *Estimated HFA* patterns for each brain-shift compensation algorithm. Specifically, we computed the negative log-transformed distance between the electrodes and the pial surface. Higher values were obtained for shorter distances and lower values were obtained for longer distances. In other words, the *Estimated HFA* patterns provide a value per electrode that indicates the intensity of the expected HFA given the electrode’s projected coordinates and allow the comparison of different back-projection algorithms.

The neuronal activity was quantified by analyzing 2 to 5 minutes of resting-state intracranial EEG activity. Patients sat comfortably in their beds and let their thoughts wander while no stimulus was presented. Channels showing epileptic activity or artifacts were removed from the analysis by visual inspection. Then, we computed the mean envelope of the HFA in the 75-145 Hz range using the Hilbert transform (Blenkmann et al., 2019). In this way, we obtained a *Measured HFA* for each electrode. Finally, we evaluated how much of the patient’s *Measured HFA* could be explained by a linear mixed-effects (LME) model with a fixed effect for Estimated *HFA* pattern and random effects for intercept grouped by electrode array and patient. We computed a LME model for each algorithm. To assess the quality of the LME models, t-tests were performed to determine the significance of fixed effects. Likelihood Ratio Tests (LRT) were used to compare models of different complexity (Bono et al., 2021), i.e., with or without random effects, with each other. Once optimal modeling was achieved, a Bayesian Information Criterion (BIC) was computed for each model (see definition in Supp. section 3.5; Schwarz, 1978). BIC is a model selection criterion that accounts for the explained variance and the number of parameters. Models with lower BIC are usually preferred.

*CEPA*’s model evidence was compared against the evidence of other models using Bayes Factor (BF). BF evaluates the ratio of the likelihood of one particular model (*i*) to the likelihood of another (*j*) and was approximated as BF = *exp* ((BIC*_i_* -BIC*_j_*)/2) (Nagin et al., 1999).

Finally, we used BF to evaluate the effect of *GridFit* vs. visually localized CT artifacts on back-projected coordinates.

## 3. RESULTS

We present two novel methods to aid the intracranial electrode localization procedure. To address the localization of CT artifacts, particularly in difficult scenarios, we propose *GridFit*, a model-based approach to automatically localize all coordinates in an electrode array. In the following subsections, we first present the use of realistically simulated scenarios to define its optimal parameters and assess its performance. Then, we validate *GridFit* with real data in challenging scenarios and compare its performance with visual procedures.

We also addressed the brain-shift compensation problem. To do this, we developed the *CEPA* method, a novel approach that combines different projection algorithms with spatial constraints. The following results compare its performance against previously validated methods in terms of error distance and modeling of electrophysiological attenuation patterns, and further expand on these results in the following subsections.

### 3.1. *GridFit* parameters definition and performance on simulated data

The *GridFit* algorithm contains many parameters that must be properly defined to obtain high localization accuracy. We estimated default parameters for the GridFit algorithm based on synthetic data provided by an open-source platform that was created for this purpose (Blenkmann et al., 2021). The parameter estimation procedure was divided into three rounds of localizing simulated CT artifacts. The first two rounds determined the best default parameters for the *GridFit* algorithm, and the third one served as a performance evaluation of the method.

During the first round, we localized a reduced set of simulated test scenarios to obtain an overview of the parameter’s effects. In particular, we noticed that *k_Corr_* and *k_Def_* had opposing effects. Increasing *k_Corr_* made the localized coordinates closer to the CT artifacts, or even caused overfitting to CT noise, while increasing the array’s deformation. On the other hand, increasing *k_Def_* produced more rigid arrays that might not follow the bending over the cortex or the curvature of depth electrode trajectories. Therefore, a correct balance between *k_Corr_* and *k_Def_* seemed fundamental for achieving precise localization results. We observed a region in the *k_Corr_* vs. *k_Def_* map space where the algorithm produces appropriate and stable results. Therefore, we set *k_Def_* to a value within this region for further steps.

During the second round, we screened over a range of *k_Corr_* values to obtain a fine-grained description of its role. To determine the optimal *k_Corr_,* we measured the normalized median error from the simulated ground truth (Eq. 10). *k_Corr_* optimal values, i.e., the ones showing smaller errors, depended on the array’s geometry (i.e., type, size, and IED) and the presence of overlaps. These optimal parameters are the ones used by the algorithm in the next sections and in the implemented version of iElectrodes (values can be found in Supp. Tables 2, 3, and 4). For more details about the results of the first two rounds of localizing simulated CT artifacts, we refer the reader to Supp. Section 2.1.

Finally, in the third round, we assessed the method’s performance. Between 16 and 60 new simulations per scenario were evaluated. We rejected 3.05% of localizations showing errors larger than 5 standard deviations for one or more contacts (183 arrays out of 6004). Overall, the accuracy was above 92% for all scenarios, i.e., localization errors below 8% of the IED, including the high noise and overlap conditions. Grids showed the best results (accuracy 95 to 99%), followed by depths (93 to 99%) and strips (92 to 99%; Figures 4A). In the most optimal conditions, i.e., low noise and no overlap, grids showed >97% accuracy, followed by depths and strips, with >95% and >94%, respectively. For practical reasons, accuracy can be easily converted into real distance errors ((1-Accuracy) * IED). For example, a 92% accuracy for a 3 mm IED grid indicates a mean error of 0.24 mm = (1-0.92) * 3 mm.

**Figure 4.**
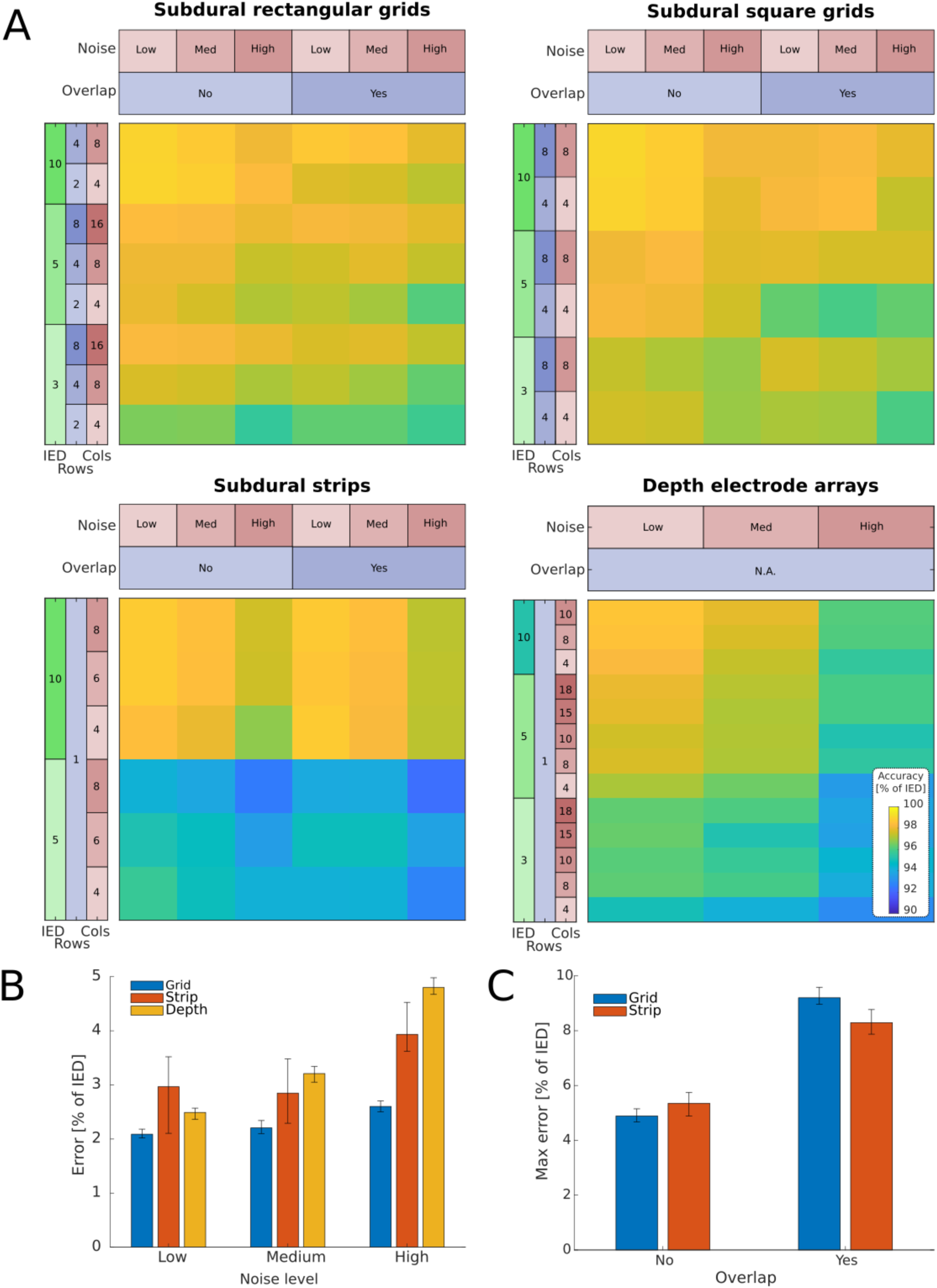
GridFit performance localizing simulated data. The GridFit algorithm performance was evaluated by localizing simulated CT artifacts under multiple realistic scenarios. Approximately 6000 simulations were used for this evaluation. **A:** Color-coded maps show the localization accuracy for rectangular and square-shaped grids, strips, and depth electrodes, with the number of rows and columns, the IED, the levels of noise, and the presence of overlaps. The color code denotes accuracy, i.e., 1 - median(dLoc Med) across arrays, where dLoc Med is the median localization error within electrodes in an array. **B** Bars show how the median error (dLoc Med, Eq. 10) increases with the noise level for the different types of arrays. ANOVA indicates medium effects of noise level and array type on the error, and a small interaction effect between them (Supp. Table 5). **C** Bars illustrate that the variation of the maximum error (dLoc Max, Eq. 11) increases in the presence of overlaps. ANOVA indicates a medium effect of overlap and a very small interaction effect between overlap and array type (Supp. Table 6). Overlaps are not available for depth arrays. Error bars denote 95% CI of the median obtained by bootstrapping. IED: Inter-Electrode Distance. Rows: Number of rows. Cols: Number of columns.

The N-way ANOVA indicated significant effects for IED, noise, overlap, array type, and noise*array type, overlap*array type, and IED * overlap interactions on the normalized median and maximum errors (Supp. Tables 5 and 6). As expected, the algorithm mean accuracy decreased with smaller IED, lower number of electrodes, lower SNR, and overlaps (Figures 4A, 4B, and 4C). ω^2^ indicates a large main effect of IED (0.23), medium effects of array type (0.07) and noise (0.09), and a small effect of overlap (0.02). Interaction effects were medium for overlap * array type (0.05), and small for noise*array type (0.02) and IED * overlap (0.02) (Lakens, 2013; Field, 2013).

Visual inspection of the results showed that the algorithm successfully localized the simulated CT artifacts in challenging situations, e.g., when an extra pair of electrodes overlap (Figure 5A) or with low SNR high-density grids (Figure 5B).

The evaluation of near-optimal parameters (plus and minus one order of magnitude in *k_Corr_*) lowered the overall accuracy by 1%. Specifically, grids’ accuracy ranged from 93 to 99%, depths’ from 91 to 98%, and strips‘ from 90 to 99%.

### 3.2. *GridFit* results from real patient data

To assess the performance of *GridFit* in real situations, we localized the CT artifacts from 20 patients implanted with grids (N = 41), strips (N = 63), and depth (N = 70) electrode arrays, and a total number of 3192 contacts. Overall, 24 arrays were implanted with overlapping electrodes from other arrays, 6 with cables or other artifacts overlapping the grids or strips, 8 with HD grids (3-4 mm IED, 1094 contacts in total), and 52 with HD depth electrodes (3.5 mm IED, 779 contacts in total).

We first examined the results qualitatively by visual inspection. *GridFit* successfully fitted the CT artifacts in all but 9 cases. Successful localizations were observed in 98% of the grids (40/41 cases), 90% of the strips (57/63 cases), and 97% of the depth electrode arrays (68/70 cases) following our criteria of all coordinates in an array lying within their respective CT artifacts.

In five of the failed automatic localizations, successful results were obtained by constraining the algorithm solutions to be aligned to a set of manually fixed coordinates at the corners or tip contacts (i.e., ***e_Fix_,*** using one, two, and four contact coordinates for depth, strips, and grid cases, respectively, 10 contacts in total). The other four unsuccessful cases were extremely curved strips (one of four and three of six contacts) where the algorithm could not converge to the CT artifacts locations using the default parameters. The manual interventions (fixed coordinates) increased the success rate to 100% of the grids, 94% of the strips, and 100% of the depth electrodes, successfully localizing 99% of the total number of electrodes (3160/3192). In the following analysis of the *GridFit* results, we considered only the correctly localized cases without manual intervention.

*GridFit* successfully localized CT artifacts in challenging situations, like cases with overlapping electrodes from other arrays (Figure 5C), connection cables running over grids (Figure 5D), high-density grids with poor SNR CT artifacts (Figure 5E), or high-density depth electrode arrays (Figure 5F).

To validate *GridFit*, we contrasted the algorithm results with the ones obtained through manual visual localization. Inferring the coordinates of some individual electrodes was difficult given the various noise sources. Overall, there was a clear consistency in the results of both approaches. Figure 5G shows the median distances (normalized by IED) between the two methods. The median distances are overall below 10% of the IED, and 6.3% for grids, 5.7% for strips, and 9.1% for depth electrodes.

To characterize the localization precision (interpreted as the spatial consistency), we computed the mean absolute deviation (MAD) of the distance between the first neighbors for each electrode array. Inter-electrode distance MAD is affected by the localization procedure’s precision and the electrode arrays’ curvature. The *GridFit* algorithm kept more uniform distances between neighbors, whereas higher variations were observed with the visual localization approach (Figure 5H, Wilcoxon signed-rank test, for grids: z = 5.49, p = 3.85 E-8; for strips: z = 6.56, p = 5.14 E-11; and for depth electrodes: z = 6.85, p = 7.09 E-12).

### 3.3. Validation of *CEPA* through pre-validated methods

To compensate for brain-shift deformations, we projected the grids (N = 40) and strips (N = 59) electrode coordinates to the SCE surface in the 15 patients implanted with them. Seven of these patients were simultaneously implanted with depth electrodes (i.e., hybrid implantations, N = 70), and therefore were also compensated for the brain deformations. The *CEPA*, *Springs*, *Normal-Grid,* and *Normal-SCE* methods were compared. The projected coordinates from the different methods showed substantial variability, as shown in Figures 6A, B, and C.

**Figure 5.**
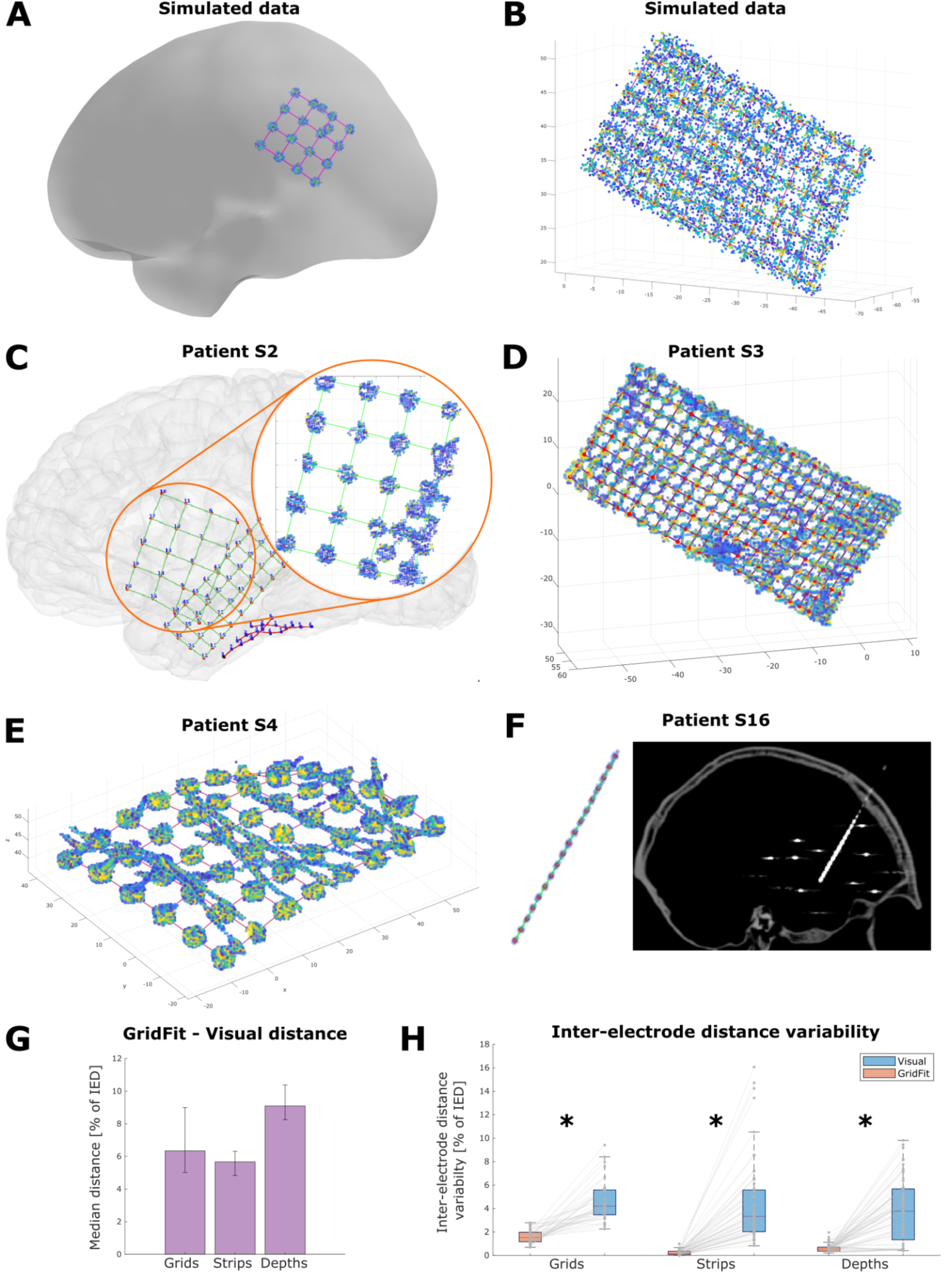
GridFit localization of CT artifacts in real and simulated data. **A.** Simulated example of overlapping electrodes. Localized CT artifacts were not affected by CT artifacts not belonging to the grid of interest (4 x 4 grid, 10 mm IED). **B.** Simulated high-density grid example with a high level of noise. The average localization accuracy in these situations is approximately 98%, i.e., 0.06 mm error (8 x 16 grid, 3 mm IED). **C.** Localization example of real overlapping grids (10 mm IED and 4 mm IED, S2). The red box shows a detail of the overlapping artifacts and the performance of GridFit in localizing the correct electrodes. **D.** Difficult localization scenario of a high-density grid with low SNR (10 x 25 grid, 3 mm IED, S3). Note that some electrodes are identifiable, whereas others are not. **E.** Localization of electrodes in a real grid with cables laying over the electrodes (6 x 8 grid, 10 mm IED, S4). **F.** Real high-density depth electrode array localized electrodes across arrays. Error bars denote 95% with low SNR (18 contacts, 3.5 mm IED, S18). The left side shows the CT artifacts and the localized coordinates (red dots), whereas the right shows the CT image. Visual localization of individual contacts in these scenarios is extremely difficult, whereas GridFit can easily handle them. **G.** Median distance between GridFit and visually localized electrodes across arrays. Error bars denote 95% CI of the median obtained by bootstrapping. **H.** Mean absolute deviation (MAD) of the inter-electrode distance (1st neighbors only) for the GridFit (orange) and visually localized (light blue) coordinates. Asterisks indicate significant differences between methods (Wilcoxon signed-rank test, p <1E-7). Points (gray) denote data from individual arrays. The center lines of each boxplot represent the median, and the edges are the 25th (Q1) and 75th (Q3) percentiles. Whiskers are located at Q1 −1.5(Q3 − Q1) and Q3 +1.5(Q3 − Q1), and outliers are plotted outside this interval. IED: Inter-Electrode Distance. SNR: Signal to Noise Ratio.

For simplicity, we present the results of back-projected *GridFit* localized coordinates in this section. The analysis of visually localized coordinates reached similar results, which can be found in the supplementary material (Supp. Section 3, Supp. Tables 7 and 8; Figures S2, S3, S4, and S5).

To assess *CEPA*’s and the other methodś performance, we first evaluated the projection error distance, i.e., the projections were contrasted against a reference location ***e_Ref_***. This reference location was obtained by averaging the results of the previously validated methods (*Springs*, *Normal-Grid*, and *Normal-SCE*). *CEPA* grid coordinates were positioned on average 0.26 mm from the reference location, with 95% of the cases at less than 0.8 mm, and at a significantly smaller distance than every other evaluated method (Figure 6D, Wilcoxon signed-rank test, z > 4 and p < 1E-4 in all cases; see Supp. Table 7 for statistical details). For strips, the distance for *CEPA* projected coordinates was smaller than 0.5 mm from the reference location and at a significantly smaller distance than the *Springs* and *Normal-SCE* methods (z > 3 and p < 1E-3; Figure S2; Supp. Table 8).

We also evaluated the error distances while considering each of the other three methods separately (*Springs*, *Normal-Grid*, or *Normal-SCE*) as a reference. *CEPA* showed better performance than the other methods in most cases (see Supp. Section 4.4, Figure S7, and Supp. Tables 12 and 13 for more details).

Moreover, we evaluated the spatial deformation roughness of the back-projected coordinates as defined by Eq. 15. Roughness is a global measure indicating how the deformations introduced by the brain-shift compensations are inconsistent. Low roughness values indicate that deformations are smooth, whereas higher values indicate a lack of spatial consistency.

Figure 6F shows the results of the pre-validated methods compared with *CEPA*. The *Springs* method depicts the smoothest results. *CEPA* showed intermediate roughness values, whereas the orthogonal-based projection methods, i.e., *Normal-Grid* and *Normal-SCE,* produced the most irregular results (see Supp. Tables 9 and 10 for statistical details).

To assess if back projections compensated for brain compressions, we evaluated the relationship between projection distance and local deformations (local IED change). *Normal-SCE, Normal-Grid,* and *CEPA* showed increased local deformation with projection distance when assessed with LME models. On the other hand, a minimal effect of distance on the local deformation was observed for the *Spring* method. Figure 6E shows the linear tendencies for each method. Additional details can be found in Supp. Section 3.3 and Supp. Table 10.

Finally, the back-projected *GridFit* coordinates were contrasted with visually back-projected ones. An overall reduction in the distance to reference and in the roughness was introduced by the use of the *GridFit* algorithm, regardless of the brain-shift compensation method used (Figures S3 and S5, Wilcoxon signed-rank test, z > 3, p < 1E-3 for projection error, and z > 4, p < 1E-5 for roughness, see Supp. Tables 7, 8, 9, and 10 for statistical details).

### 3.4. Validation of *CEPA* using electrophysiological data

To independently quantify the quality of the different back-projection algorithms, we compared the explanatory power of their respective *Estimated HFA* predicting resting-state *Measured HFA* (Figure 7A-C). The *Estimated HFA* patterns reflect the expected HFA solely based on the distance of the projected coordinates to the cortical surface (Figure 7B). The intensity of the recorded neuronal activity was expected to be larger for electrodes located over gyri (closer to the cortex) and smaller for electrodes located over sulci (further from the cortex).

**Figure 6.**
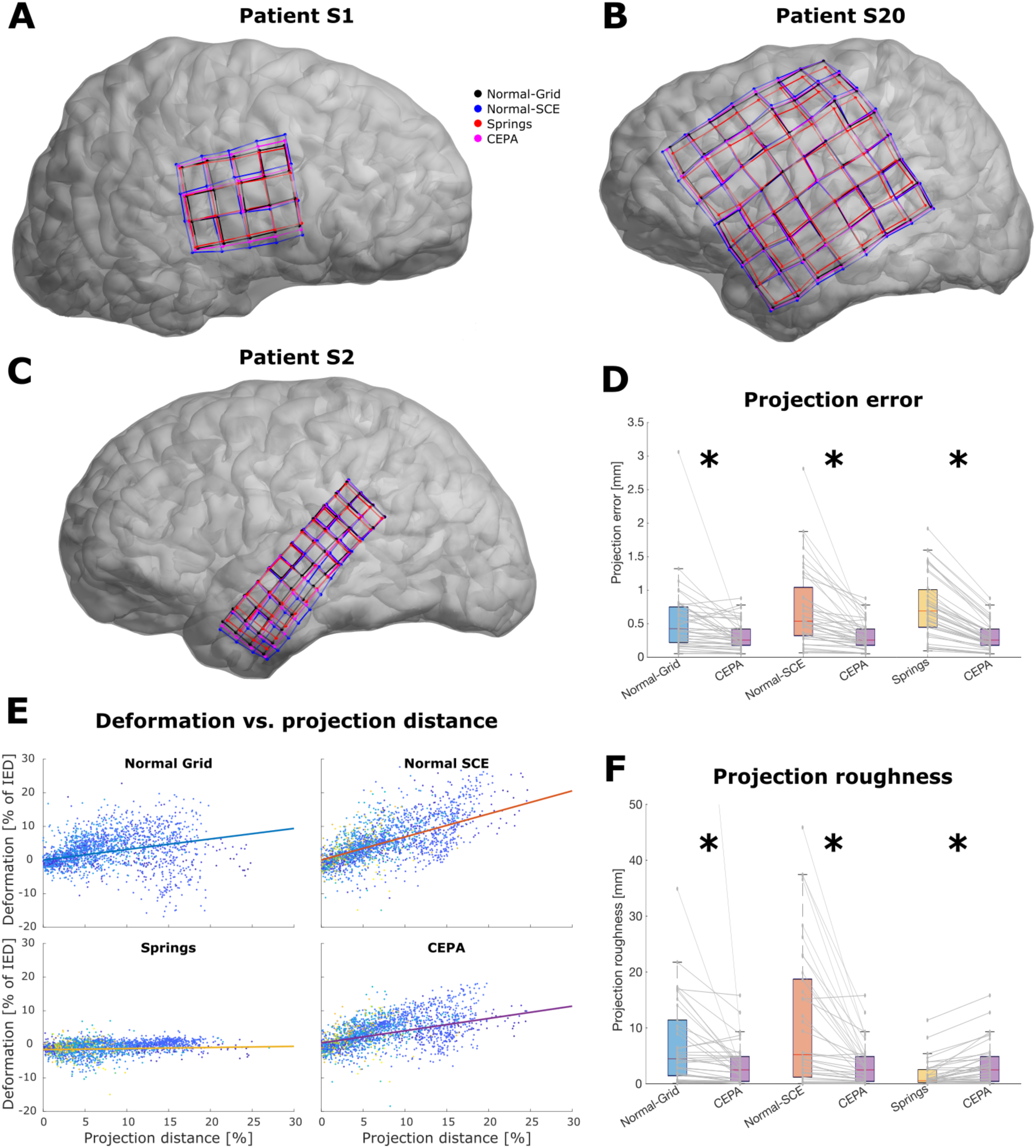
CEPA brain-shift compensation compared to other methods. **A, B**, and **C**: Plots show the behavior of brain-shift compensation algorithms in three examples (S1, S20, and S2, respectively). Grids of different sizes and geometries were back-projected using Normal-Grid (black), Normal-SCE (blue), Springs (red), and CEPA (magenta) methods. Results differed substantially, and Normal-SCE projections were typically the most expanded, whereas Springs projections were the least expanded. In comparison, Normal-Grid and CEPA typically showed moderate levels of expansion. **D**: Boxplots show the projection errors for the different brain-shift compensation algorithms contrasted against CEPA. The projection error for CEPA was significantly smaller than for all other methods. Points (gray) denote data from individual grids. **E.** Deformation vs. projection distance for different projection methods. Deformation was computed relative to the IED, whereas projection distance was computed relative to the distance to the brain’s center of mass. Each point represents an electrode color-coded by patient, and the lines depict the tendency obtained via linear mixed-effect models. In contrast to the other approaches, the Springs method was not sensitive to changes in the projection distance, which is consistent with the almost null roughness observed. **F**: Boxplots show the projection roughness of the different algorithms contrasted against CEPA. Lower values indicate smoother or more regular spacing in the projected coordinates. CEPA showed significantly smaller roughness than the Normal-Grid and Normal-SCE approaches and higher roughness than the Springs method. Points (gray) denote individual grids. For illustrative purposes, a single point for the Normal-Grid method is not shown. The center lines of each boxplot represent the median, and the edges are the 25th (Q1) and 75th (Q3) percentiles. Whiskers are located at Q1 −1.5(Q3 − Q1) and Q3 +1.5(Q3 − Q1), and outliers are plotted outside this interval. Asterisks indicate significant differences between methods (p < 1E-4, Wilcoxon signed-rank test). IED: Inter-Electrode Distance

**Figure 7.**
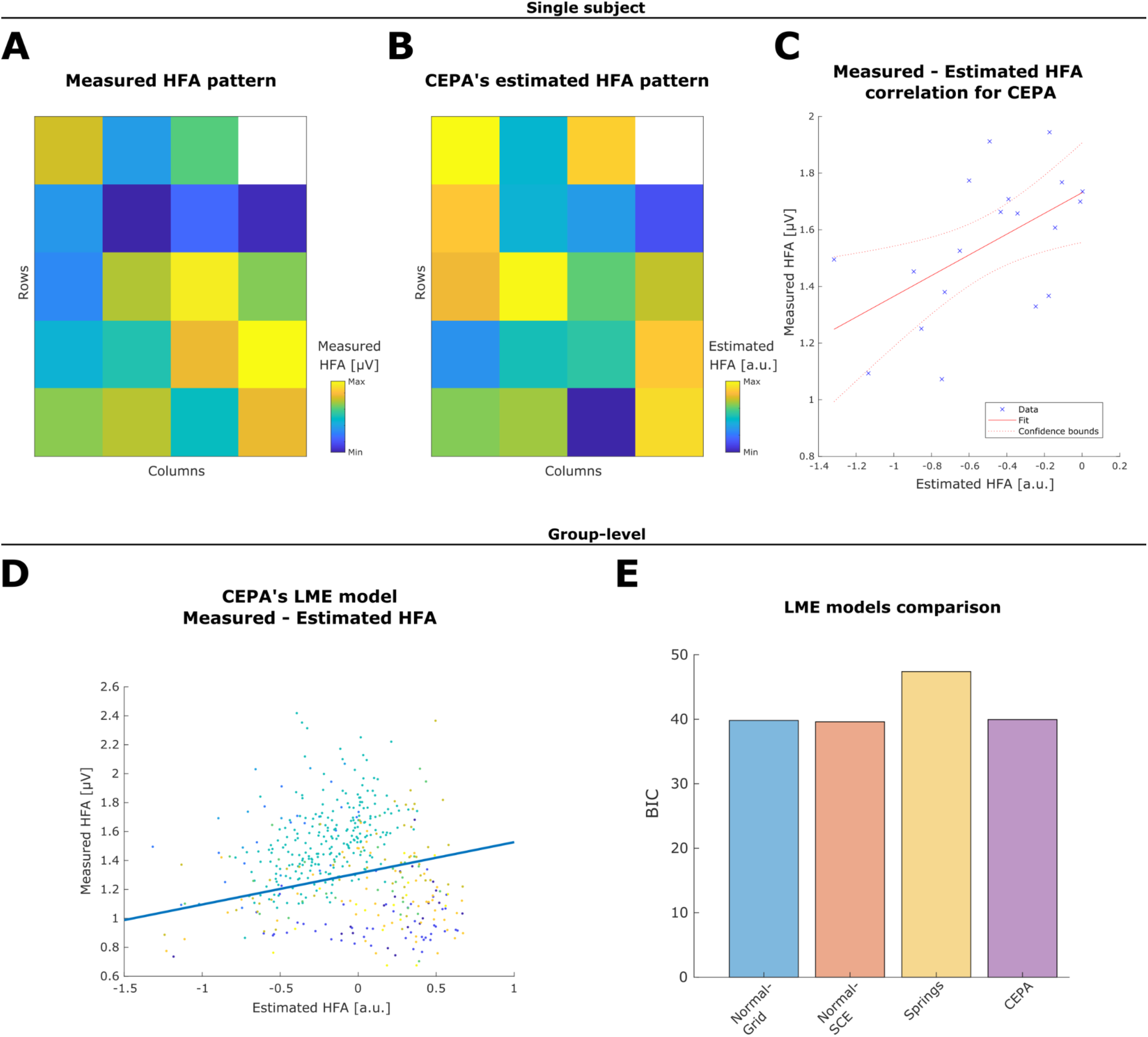
HFA resting-state data and HFA estimation. **A.** Measured HFA mean value during resting state from an example 5 x 4 grid (Patient S1). **B.** Estimated HFA pattern computed from back-projected CEPA coordinates of the same grid (Patient S1). The larger the distance of an electrode to the cortex, the smaller the Estimated HFA (stronger attenuation). **C.** Correlation between the CEPA’s HFA estimated pattern and the measured HFA as shown in A and B. **D.** Group-level (N = 5 patients) linear relationship between the measured HFA and the estimated HFA for CEPA back-projected coordinates. Individual dots represent single electrode data, color-coded by patient. **E.** Bayesian Information Criterion (BIC) compares the different LME models of the respective back-propagation methods. Lower BIC values indicate that more data variance is explained by the model (better fitting).

Respective *Estimated HFA* patterns were computed for each brain-shift correction algorithm in nine grids from five patients. We used LME models to estimate on a group level to what extent the *Measured HFA* is predicted by the *Estimated HFA* (Figure 7D). The explained variance ranged from 46.67 to 47.55%, as estimated by the adjusted R^2^. *Springs* showed the lowest explained variance, while the other three methods showed similarly higher values, indicating more explanatory power.

The LME model comparison via BIC analysis indicates similar performance for *CEPA, Normal-SCE,* and *Normal-Grid* (Figure 7E). The *Estimated HFA* pattern obtained from these methods’ coordinates explained a similar amount of information in the recorded resting-state HFA. The *Springs* method showed a larger BIC, indicating an inferior performance compared to the other methods and therefore being the least preferred alternative.

The likelihood ratio of two competing hypotheses, given by the Bayes Factor, indicates strong positive evidence for the use of *CEPA* versus the *Springs* method (BF odds ratio 41.32), whereas the evidence for choosing between *CEPA* versus *Normal-Grid* or *Normal-SCE* was insensitive (BF odds ratio 0.93 and 0.84 respectively; Kass and Raftery, 1995). Supp. Table 14 shows a summary of the LME model results.

Additionally, we incorporated the evidence from all LME models to evaluate the impact of CT artifact localization strategies on the back-projection coordinates. BF provided decisive positive evidence for using *GridFit* versus visually localized coordinates (BF odds ratio 9,520; Supp. Table 15 shows the LME model results for visually localized coordinates).

## 4. DISCUSSION

Accurate anatomical registration of intracranial electrodes is essential for precisely interpreting findings in clinical and cognitive neuroscience iEEG studies. Localization accuracy is affected by: i) the procedure used to locate the post-implantation CT artifacts; and ii) the procedure chosen to compensate for brain shifts in grid and strip cases. Both impact the accuracy of the final results, and errors in the first procedure can propagate to the second. To address these issues, we introduced two novel algorithms for the anatomical registration of intracranially implanted electrodes. First, we presented the *GridFit* algorithm to localize CT artifacts and assessed its accuracy with both real and realistically simulated data. We demonstrate that the method excelled in challenging scenarios with real data and provided more precise electrode locations than visual procedures. Results were highly reliable, even in low SNR or high-density array cases.

Second, we introduced *CEPA* to back-project implanted grid and strip coordinates to the cortical surface to compensate for brain shifts. Our analysis indicates that the novel algorithm provided projections to the cortical surface envelope that were accurate, spatially smooth, increased IED with the projection distance, and predicted electrophysiological activity patterns.

In the following, we discuss these results in more detail and compare our methods with other available alternatives. Finally, we consider the limitations of our study and remaining challenges.

### 4.1. *GridFit* algorithm for the localization of CT artifacts

Low SNR CT artifacts are especially challenging for localization algorithms. Low SNR occurs in the presence of metallic objects in the proximity of the electrodes, overlapping electrodes, or high-density electrodes (HD arrays). Low spatial image resolution or unrelated artifacts can preclude automatic algorithms from detecting individual contacts. Manual interventions are often required to identify these lost contacts (Branco et al., 2018a; LaPlante et al., 2016; Taimouri et al., 2014), and arrays with overlapping electrodes might need to be excluded from the analysis (Brang et al., 2016). Moreover, the presence of electrodes in the proximity of the ones of interest has been associated with localization errors (Narizzano et al., 2017), while noise in CT images has been mistakenly detected as electrodes (La Plante et al., 2016).

With the *GridFit* method, we introduce a novel approach to locate CT artifacts developed to address these challenging scenarios. *GridFit* simultaneously localizes all the electrodes in grids, strips, or depth arrays using flexible models fitted to the CT artifacts.

We tested our method with an extensive database of realistically simulated scenarios where the exact locations are known. Some of these scenarios presented high levels of noise and overlapping artifacts (Figure 5). Nevertheless, the method succeeded in locating the electrode coordinates with high accuracy. The accuracy was above 92% in all cases, i.e., errors below 8% of the IED (Figure 4).

Overall, we observed a tendency to obtain higher localization accuracy and precision with lower noise levels, a higher amount of electrodes that could be localized, a larger IED, and the absence of electrode overlaps. In sum, localizing electrodes in simulated scenarios indicates that the higher the quality or the quantity of the data, the better the localization. Notably, the algorithm’s accuracy was lowered by only 1% when selecting suboptimal parameters, indicating the procedure’s reliability. **[real data]** We further validated *GridFit* with 174 real electrode arrays obtained from 20 patients (3192 contacts), where localization difficulties were a prerequisite for inclusion in this study.

Patients had either low SNR images, grids underneath cables, overlapping grids, presence of artifacts, or HD arrays (IED ≤ 4mm), as shown in Figure 5. These characteristics rendered the testing dataset challenging (Centracchio et al., 2021; Branco et al., 2018a). Despite such obstacles, the *GridFit* algorithm automatically localized 95% of the arrays successfully. By manually adding fixed coordinates in 5 of the most difficult cases (one to four coordinates per case), we achieved an overall 98% successful localization of arrays, and 99% of the contacts. In four highly curved, C-shaped strips (22 contacts in total), we could not achieve successful localizations, revealing limitations of the proposed method. Ad-hoc deformation parameters proved helpful in addressing this specific problem but were not further analyzed given the small number of cases observed.

Comparably, Davis et al. (2021) reported a 93% sensitivity in the automatic detection of electrodes, and Centraccio et al. (2021) reported between 88 to 99% classification sensitivity using geometrical features extracted from CT artifacts of metallic objects. Moreover, an exceptional 99% accuracy was obtained in distinguishing between electrodes and other metallic objects. Such automatic classification approaches could easily be combined within *GridFit* to offer even more reliable methods. For example, localized coordinates could be integrated as anchors weighted according to the reliability of the predictors (Nicora et al., 2022).

Localization of HD arrays, where the IED is ≤ 4mm, can be of particular challenge. Branco et al. (2018a) correctly localized all the 4 mm grid contacts using their automatic algorithm (3 cases), but most of the contacts had to be manually localized in the 3 mm cases (2 arrays). Interestingly, the localization of HD depth arrays has shown higher detection rates, probably due to their constrained geometry and less likely deformations. Davis et al. (2021) reported an 87% sensitivity in the automatic detection of HD depth contacts, while Narizzano et al. (2017) reported 87% utilizing the planned implantation target and surface entry coordinates, and up to 99% when the tip or entry point coordinates were manually provided. In comparison, *GridFit* successfully localized the CT artifacts in all 60 HD arrays tested. Eight of these cases were grids (1094 contacts in total), and the other 52 were depth arrays (779 contacts in total). Only two arrays (one grid and one depth) required manual intervention by visually localizing corner or tip contacts. Such a procedure was embedded in the iElectrodes GUI, making reliable localizations straightforward.

We compared *GridFit* with typical visually localized coordinates. The distances between the approaches were 10% of the IED, similar to the results reported by Blenkmann et al. (2017) for grids and depth electrodes with 10 and 5 mm IED. Visually localizing the coordinates without technological aids has an intrinsic mean observer error of approximately 0.8 mm in high SNR situations (Blenkmann et al., 2017). This error is presumably larger in challenging scenarios where the noise in the images or other artifacts make the procedure even harder, impacting the quality of the results and increasing the procedure’s duration (Narizzano et al., 2017). Therefore, visually localized coordinates should be considered a reference, not a gold standard.

A validation and comparison between *GridFit* and the visual approach was provided by measuring the deformations introduced in the IED, since the IEDs are expected to be minimally affected by the bending of the arrays (<0.5%, Blenkmann et al., 2021). Electrode coordinates defined by *GridFit* were smoother (smaller deviations of the IED) than the ones obtained by visual inspection, demonstrating that the *GridFit* results are more reliable than the visual ones.

Overall, the model-based *GridFit* algorithm showed highly reliable results with both real and simulated data. Notably, performance excelled in challenging scenarios, such as low SNR or high-density array cases. The excellent performance of *GridFit* is likely due to the model-based approach to localize all contacts simultaneously. The subset of image voxels under analysis might have missing or misleading information about individual electrode coordinates, but as a whole, they provide sufficient information to accurately and reliably localize all contacts in an array.

### 4.2. *CEPA* for brain-shift compensation of subdural grids and strips

Brain shifts up to 10-20 mm are common after implanting subdural grids or strips (LaViolette et al., 2011; Studholme et al., 2001). Such brain deformation constitutes a complex problem involving multiple variables, such as the size and location of the skull opening, the head orientation during surgery, the amount of cerebrospinal fluid lost and reabsorbed, and the swelling of soft tissue (Roberts et al., 1998; Studholme et al., 2001). These deformations need to be accurately corrected to associate electrophysiological activity with the underlying anatomy with high spatial resolution.

Post-implantation brain-shift deformations preclude accurate localization directly from intracranial photographs or post-implantation CT images. Acknowledging the implications of these deformations in reconstructing the electrode locations on the pre-implantation brain scans is essential. The issue is usually overlooked despite affecting post-implantation CT and photography-based corrections. Remarkably, using a simple formalization of the electrode back-projection problem, Brang and colleagues (2016) revealed that the array shape and IED are not guaranteed to be preserved in the back-projected coordinates. For example, in a typical implantation surgery, a grid is placed over the surface of a non-linearly deformed brain, but crucially, the grid shape and its IED are preserved. Therefore, array coordinates projected to the non-deformed pre-implantation MRI brain surface are expected to compensate for this behavior, typically expanding the arraýs surface. Notoriously, expansions are limited in back-projection approaches that penalize array deformations, like the *Spring mesh* family of methods (Dykstra et al., 2012; Trotta et al., 2018). In other words, since IED deformations are penalized, projections only suffer small IED changes (Figure 6F) and are unaffected by the projection distance (Figure 6E).

On the other hand, the *Orthogonal projection* family of methods (Hermes et al., 2010; Kubanek & Schalk, 2015; Brang et al., 2016) computes projections at the individual electrode level. Projections tend to increase the IED proportionally to the projection distance (Figure 6E), which is theoretically expected to compensate for brain compression. However, these methods produce less smoothly deformed solutions, i.e., spatially contiguous changes in the IED are more heterogeneous (Figure 6F), because orthogonal projection vectors are computed individually for each electrode.

Considering the spatially continuous brain-shift deformation, a certain degree of smoothness in the array’s deformations is anticipated. Given the previously described inaccuracies and limitations, we combined both approaches in *CEPA* to get the best of each family of methods: Back-projections following orthogonal orientations, while keeping spatially smooth variations in the IED.

To assess *CEPA’*s performance, we contrasted it with previously validated methods from the *Spring mesh* and *Orthogonal projection* families (Dykstra et al., 2012; Trotta et al., 2018; Hermes et al., 2010; Kubanek & Schalk, 2015; Brang et al., 2016). These methods were validated against photography-based approaches (2-3 mm error, Dykstra et al., 2012; Hermes et al., 2010; Branco et al., 2018a). Results were highly dissimilar between methods, as shown in Figure 6. *CEPA* outperformed the other methods by producing locations significantly closer to the reference (less than 0.3 mm error, Figure 6D). As expected, *CEPA* estimations were also smoother, and the projection distance effects on deformation were less prominent than in the orthogonal projection methods (Figures 6E and 6F). The opposite effect was observed when compared with the *Springs* method. Therefore, *CEPA* achieved solutions that retained the benefits of both back-projection families.

In this study, we introduced *Estimated HFA* patterns as a performance marker for localization methods. HFA is generated within a few millimeters of the recording electrodes and mirrors the average spiking activity of adjacent neurons (Ray & Maunsell, 2011; Mc Carty, 2021, Leszczynski et al., 2020), and is significantly attenuated as a function of the distance between the electrodes and the underlying cortex. Cerebral veins naturally lie in sulcal folds, and the presence of blood vessels underneath electrodes can dampen the recorded signals by 30 to 40% (Bleichner et al., 2011).

Computing the correlation between *Measured HFA*, derived from resting-state recordings, and *Estimated HFA,* based on the distance between electrodes and cortex, has been shown as a reliable approach to localize HD grids (Branco et al., 2018b). Given the lack of a gold standard for localized coordinates, modeling HFA patterns provides an alternative and independent measure that is simple to obtain, theoretically sound, and empirically closer to the primary aim of intracranial EEG, which is the assessment of electrophysiological activity.

In our study, we modeled HFA patterns to compare the performance of different back-projection methods. Patterns obtained from *CEPA, Normal-Grid,* and *Normal-SCE* back-projected coordinates explained the *Measured HFA* variance substantially better than the *Springs* method. BF model comparison provided decisive evidence for using *CEPA* against *Springs*, but indecisive evidence when compared with orthogonal projection methods. Importantly, these methods also showed increased deformations (expansion) associated with the projection distance, in contrast to the minimally deforming *Springs* method (Figure 6E). This association suggests that the expansion of arrays is an essential feature for back-projection algorithms compensating for brain compressions.

When the localization and back-projection results are considered together, they indicate that combining *GridFit* and *CEPA* provides the best outcome. The localization accuracy was higher than with any other combination of methods, and the HFA modeling based on *GridFit* and *CEPA* was among the best-performing alternatives with respect to the explained variance. Interestingly, our observations based on localization accuracy and HFA modeling supported the use of *GridFit* and improved the back-projection outcome. This indicates that errors propagate from the CT artifact localization step to the brain-shift compensation step. Finally, the back-projection results can differ substantially between methods. Projected coordinates can “jump” from one cortical gyrus to the next, depending on the algorithm used. For this reason, we recommend that users consider this uncertainty when interpreting normal or pathological electrophysiological activity.

### 4.3. Limitations and challenges

Although *CEPA* and *GridFit* proved to be successful tools, some limitations should be noted.

i) We used a predefined and fixed set of parameters in the *CEPA* implementation. Although the high performance obtained with these parameters, it is likely that other (i.e., optimal) values offer even better performance. Given the relatively small number of cases, searching for optimal parameters was not possible. A promising approach in this direction is the use of simulated data (e.g., following Blenkmann et al., 2021).
ii) Our back-projection approach was contrasted with a restricted number of methods. This approach is not an extensive evaluation of all possible methods but a comparison with some of the most commonly used and representative techniques. Error distance to an average location was used to compare *CEPA* with the other methods. Although the choice of reference rests on a parsimonious assumption, it is difficult to argue that the *reference* location is the best one. Nevertheless, when considering the situation that any other method produces the best solution, *CEPA* was either a significantly better alternative or performed equally to the others.
iii) The modeling of HFA has limitations. The correlation between *Measured HFA* and *Estimated HFA* patterns has been used to quantify the performance of localization methods. However, we employed resting-state iEEG data from a small group of patients. We were not able to measure differences between *CEPA* and the *Orthogonal projection* family methods, probably because of the functional brain activity variability of the participants. For example, internally and externally directed attention might engage distinct functional neural networks and activity patterns (Kam et al., 2019), which might be reflected in the *Measured HFA* patterns, and therefore affect the performance evaluation procedure.

Novel reference structures were recently developed to allow post-implantation MRI-based localization of microelectrodes, avoiding high doses of ionizing radiation exposure incurred by patients during CT scanning (Erhardt et al., 2020). These technologies use MRI-visible patterns that might benefit from model-based approaches, like the one used in *GridFit*, to provide accurate localizations in real patient implantations.

### 4.4. Open-source and GUI availability

The methods described in this article are publicly available on the iElectrodes site (https://sourceforge.net/projects/ielectrodes/), as Matlab® scripts. iElectrodes is a popular open-source toolbox for intracranial electrode localization, with more than 1700 downloads to date. Its graphical user interface (GUI) aids visualization and user interaction, data management, generation of reports, and sharing of localization cases. Data import-export functionalities allow simple integration with other popular analysis toolboxes (e.g., Fieldtrip, EEGLAB) and brain atlases, among other valuable tools.

## 5. CONCLUSIONS

We introduced two new methods for the localization of intracranial grid, strip, and depth electrodes. First, we developed the *GridFit* algorithm to achieve robust localization of electrodes from CT images. We tested our algorithm with challenging cases where other commonly used methods have documented suboptimal performance or failure. *GridFit* produced highly accurate results even in the case of low SNR CT images, cases with overlapping cables or other artifacts, and high-density array cases. It outperformed electrode localization via visual inspection, which is still a common practice in this field. Extensive simulations and real data supported these conclusions.

Second, we developed *CEPA* to enable smooth and orthogonally expanding back projections during the brain-shift compensation of grids and strips. Compared to other available methods, *CEPA* showed the highest spatial precision (i.e., the slightest error distance) and was among the top performers in predicting electrophysiological activity. Altogether, *GridFit* and *CEPA* showed high-accuracy anatomical localization of implanted electrodes, even in the most challenging implantation scenarios. Moreover, both algorithms are implemented in the iElectrodes open-source toolbox, making their use simple, accessible, and easy to integrate with other popular toolboxes used for analyzing electrophysiological data.

## 6. DECLARATIONS

### 6.1. Ethics approval and consent to participate

Patients were recruited from Albany Medical College; University of California, Irvine; University of California, San Francisco; and Oslo University Hospital.

This study was approved by the Regional Committees for Medical and Health Research Ethics, Region North Norway (REK 2015/175), and the Human Subjects Committees at UCSF, UC Irvine, UC Berkeley, and Albany Medical College, in accordance with the ethical standards laid down in the 1964 Declaration of Helsinki.

All patients gave their written informed consent for participation in this study and the use of their collected information.

### 6.2. Availability of data

The patients’ datasets analyzed during the current study are not publicly available due to our ethical approval conditions that do not permit the public archiving of anonymized study data.

## 6.3 Acknowledgments

We are grateful to the patients for kindly participating in our study.

Special thanks to Dr. Ludovic Bellier for the critical discussions that triggered this project. We appreciate FRONT Neurolab - RITMO, LEICI, and ENyS members for their rich discussions and feedback. Thanks to Santiago Collavini and Julian Fuhrer for their useful comments on the manuscript. We also acknowledge Drs. Julia Kam and Ludovic Bellier for providing the resting-state data.

The simulations in this project were performed on resources provided by UNINETT Sigma2 - the National Infrastructure for High-Performance Computing and Data Storage in Norway.

This work was partly supported by the Research Council of Norway, project numbers 240389 and 314925, through its Centres of Excellence scheme, project number 262762, and Intpart project number 274996, NINDS Grant R37NS21135, NIMH CONTE Center P50MH109429, and Brain Initiative U01-NS108916 and U19NS107609.

## 6.4. Authors’ contributions

Conceptualization: AOB, TE, and AKS. Data curation: TE, JL, EC, PB, GS, JI, PGL, RTK, AKS. Software development, investigation, methodology: AOB. Funding acquisition: AOB and AKS. Project administration: AOB, TE, and AKS. Validation of results: All authors. Writing - original draft: AOB. Writing - review & editing: All authors. All authors read and approved the final manuscript.

## 6.4. Competing interests

The authors declare no competing interests.

## List of abbreviations

BIC: Bayesian Information Criteria
BF: Bayes Factor
CEPA: Combined Electrode Projection Algorithm
ECoG: Electrocorticography
EEG: Electroencephalography
MRI: magnetic resonance imaging
fMRI: functional MRI
HFA: High-Frequency Activity
HD: High-Density
IED: Inter-Electrode Distance
iEEG: Intracranial EEG
LME: Linear Mixed-Effects
LRT: Likelihood Ratio Tests
MAD: Mean Absolute Deviation
PCA: Principal Component Analysis
SEEG: Stereo EEG
SCE: Smooth Cortical Envelope
SNR: Signal-to-Noise Ratio

## Equation’s variables and constants

*d_o jk_* initial distance between points *j* and *k*

*d_Lo_*_c *Max*_ maximum normalized localization error

*d_Loc Med_* median normalized localization error

*d_jk_* distance between points *j* and *k*

*d_Proj j_* normalized projection distance for electrode *j*

***e_0_*** initial electrode locations

***e_A_^S^*** anchored coordinates from Normal-SCE projection

***e_A_^G^*** anchored coordinates from Normal-Grid projection

***e*** electrodes location

***e_GridFit_*** *GridFit* localized coordinates

***e_Vis_*** visually localized coordinates

*k_Corr_* co-registration weighting constant

*k_Def_* deformation weighting constant

*k_Trans_* translation weighting constant

*k_Anch_* anchoring weighting constant

*k_Rough_* roughness weighting constant

***m*** structural model coordinates

***t****_k_* back-projection displacement vector for electrode *k*

***v*** voxel coordinates

*w* voxel intensities

*D* inter-electrode distance

*E_Corr_* co-registration cost

*E_Def_* deformation cost

*E_Trans_* translation cost

*E_Anch_* anchoring cost

*E_Rough_* roughness cost

***G_H_*** horizontal deformation matrix

***G_V_*** vertical deformation matrix

***K*** nine-point stencil kernel

***L_H_*** horizontal Laplacian matrix

***L_V_*** vertical Laplacian matrix

*N_Vox_ (N)* number of thresholded voxels

*N_Elec_ (P)* number of electrodes

*N_Conn_ (L_e)_* number of connections within a model

*N_Adj_ (C)* number of adjacent pair of electrodes

*N_Anch_* number of anchoring methods

*N_Rows_* (A) rows

*N_Cols_* (B) columns

*N_Mod_ (M)* number of structural model points

*T* array thickness

σ spatial dispersion constant of the co-registration

### Notes

Lowercase bold indicates vectors.

Uppercase bold indicates matrices.

## Notes

### Competing Interest Statement

The authors have declared no competing interest.

### Summary of Updates

minor typos in the abstract

