## Supplementary material for "Anatomical registration of intracranial electrodes. Robust model-based localization and deformable smooth brain-shift compensation methods"

#### 1. Patient overview

Supplementary Table 1

| N | Age | Sex | Electrode Arrays | IED | HD | Array type | Hem | Lobe | Overlaps | Comments |
|---|---|---|---|---|---|---|---|---|---|---|
| 1 | 54 | M | 4x5, 3x4, 1x4, 4x8, 1x4, 1x4 | 7, 10 | | G, S | R | F, T, P, O | G4-6 | Over resected brain. Big artifact on top (G4) |
| 2 | 19 | F | 1x4, 1x6, 1x6, 1x6, 4x5, 4x12 | 6, 10 | | G, S | L | F, T, P | G5-6 | Highly curved (S3) |
| 3 | 33 | F | 1x4, 1x4, 1x4, 1x4, 1x4 / 1x6, 2x3, 1x4, 1x4, 1x4, 10x25 | 10 / 10, 3 | G11 | S / S, G | L / R | F, T, P / F, T | G9-10 | Highly curved (S3-4, S10)<br>Low SNR (G11)<br>Fixed coordinates added (S4, G11) |
| 4 | 57 | F | 1x4, 1x6, 1x6, 1x4, 1x4, 6x8, 1x6, 1x6, 1x6, 4x12 | 6, 10 | | G, S | R | F, T, P | | Cables on top (G6, S7)<br>Noisy artifacts (G10) |
| 5 | 31 | M | 1x4, 1x4, 1x4, 1x4, 1x4 / 4x12, 2x8, 1x4, 1x4, 1x4, 2x4, 2x6, 2x3, 1x4, 1x4, 1x4 | 6, 10 | | S / S, G, D | R / L | T, P / F, T, P | D8-9, G11-12 | |
| 6 | 33 | M | 1x4, 1x4, 2x5, 1x4, 1x4, 1x4, 2x10, 2x17, 6x25 | 3, 10 | G7-9 | S, G, D | L | F, T, P | S4-5 | Extremely curved - failed GridFit localization (S5)<br>Highly curved (S6)<br>Cut grid (G7-9) |
| 7 | 21 | M | 1x4,1x4, 1x4, 1x4, 4x5, 1x6, 1x6, 1x6, 16x16 | 10, 4 | G9 | G, S | R | F, T, C | | Highly curved (S6-8, G9)<br>Fixed coordinates added (S8)<br>Artifacts on top. Low SNR (S9) |
| 8 | 23 | M | 4x5, 1x6, 1x6, 1x6, 1x4, 1x4, 1x4, 1x8, 16x16 | 4, 5, 10 | G9 | S, G, D | R | F, T, P, O | | Highly curved (S2-4) |
| 9 | 52 | F | 1x10, 1x10, 1x6, 1x6, 1x6, 1x4, 1x4, 1x4, 8x8 | 5, 10 | | G, S, D | L | F, T, P, O | | Abnormal temporal lobe (resection cavity)<br>Highly curved (D1)<br>Extremely curved - failed GridFit localization (S3-5) |
| 10 | 21 | M | 1x8, , 1x8, 1x8, 1x8 / 1x9, 1x8, 1x8, 4x8, 8x8 | 5, 10 | | D / G, D | R / L | T / F, T, O | G8-9 | |
| 11 | 26 | F | 8x8, 4x5, 8x8 | 4, 10 | G3 | G | L | F, T, P | | Highly curved (G1-2)<br>Low SNR (G3) |
| 12 | 33 | F | 4x8, 4x4, 4x4, 8x8, 1x6, 1x7, 1x6, 1x6, 1x10, 1x10, 1x10 | 4, 5, 10 | G4 | G, S, D | L | F, T, P | G1,D11 | Highly curved (S5-8, D9-11) |

| | | | | | | | | | | |
|---|---|---|---|---|---|---|---|---|---|---|
| 13 | 20 | F | 4x8 ,4x4, 2x5, 8x8 | 10 | | G | L | F, T, P, O | G3-4 | Temporal resection<br>Highly curved (G4) |
| 14 | 41 | F | 1x10, 1x17, 1x15, 1x12, 1x12, 1x15, 1x15, 1x15 | 3.5 | D1-8 | D | R | F, P | | |
| 15 | 35 | M | 1x18, 1x18, 1x15, 1x15, 1x15, 1x15, 1x12, 1x12, 1x16, 1x15 | 3.5 | D1-10 | D | L | F, P | | |
| 16 | 28 | M | 1x15, 1x12, 1x12, 1x9, 1x8, 1x8, 1x9, 1x12, 1x12, 1x12, 1x9, 1x11, 1x11, 1x11 | 3.5 | D1-14 | D | L | F, T, P, I | | Low SNR. Fixed coordinates added (D1) |
| 17 | 23 | M | 1x17, 1x18, 1x18, 1x12, 1x15, 1x15, 1x15, 1x15 | 3.5 | D1-8 | D | L | P | | |
| 18 | 38 | M | 1x18, 1x18, 1x8, 1x10, 1x10, 1x10, 1x15, 1x15, 1x15, 1x15, 1x15, 1x15 | 3.5 | D1-12 | D | L | F, T, I | | Low SNR (D1-2) |
| 19 | 26 | F | 1x6, 1x6, 4x5, 12x4, 1x6, 1x6, 1x6 | 6, 10 | | G, S | L | F, T, P | | Artifacts on top. Low SNR (G4) |
| 20 | 48 | F | 8x8, 4x4, 2x6, 1x10, 1x10, 1x10 | 5, 10 | | G, D | L | F, T, P | G1-2, D4-6 | Cables on top (G1-2)<br>Curved. Fixed coordinates added (D6) |

IED: Inter-electrode distance. HD: High-density (IED $\leq$ 4mm). Hem: Hemisphere.
G: Grid, S: Strip, D: Depth. L: Left. R: Right. F: Frontal. T: Temporal. P: Parietal. O: Occipital. I: Insula. C: Cingulate.
/: separate left and right implants.

#### 2. *GridFit* algorithm supplementary material

##### 2.1. Definition of GridFit parameters using simulations

Three rounds of localizing simulated CT artifacts was used to determine the best parameters for the *GridFit* algorithm. The first round was intended to get an overview of the effects of the parameters, the second round to have a fine-grained description of the parameters, and the third one to assess the method's performance.

After the first round, we noticed that $k_{Corr}$ and $k_{Def}$ had opposing effects. For the second round, we defined all parameters for the first fitting step, i.e., $k_{Trans}$= 1E-6, $k_{Def}$= 1E5, $k_{Corr}$ = 1E5, σ = IED, ε = 1 [mm], and defined $k_{Trans}$ = 1E-2, $k_{Def}$ = 1E5, and σ= ¼ IED for the second fitting. We evaluated the algorithm's performance by varying the $k_{Corr}$ parameter ($k_{Corr}$ range 10 to 1 E11, localizing in total ~69000 simulated cases). We ran the second fitting step without constraints to incorporate a larger range of parameters with less computational time (Eq. 7, 8, and 9). We observed that optimal $k_{Corr}$ parameters, i.e., the ones producing a smaller normalized median error (Eq. 10) or higher accuracy, depended mostly on: i) electrode type, ii) the number of electrodes, iii) the IED, and iv) the presence of overlaps (for grids and strips). Smaller variations were observed to changes in noise levels or local curvature. Therefore, we defined a function that automatically computes the optimal $k_{Corr}$ parameter, i.e., $k_{Corr\ Opt}$, considering the four variables. Supplementary Tables 2, 3, and 4 show the optimal values obtained for each case. Optimal $K_{Corr}$ value for other geometries than the evaluated ones are obtained by a modified Akima cubic Hermite interpolation (using the *interp3* function in Matlab).

Supplementary Table 2. $k_{Corr\ Opt}$ values for Grids

| Grid | No overlap | | | | | Overlap | | | | |
|---|---|---|---|---|---|---|---|---|---|---|
| IED[mm]\ N ch | 8 | 16 | 32 | 64 | 128 | 8 | 16 | 32 | 64 | 128 |
| 3 | 1E6 | 1E6 | 1E7 | 1E8 | 1E8 | 1E5 | 1E6 | 1E7 | 1E7 | 1E8 |
| 5 | 1E6 | 1E7 | 1E7 | 1E8 | 1E9 | 1E6 | 1E6 | 1E7 | 1E8 | 1E9 |
| 10 | 1E7 | 1E9 | 1E9 | 1E10 | 1E10 | 1E6 | 1E7 | 1E8 | 1E8 | 1E9 |

N ch = number of electrodes; IED = Inter-electrode distance

Supplementary Table 3. $k_{Corr\ Opt}$ values for Strips

| Strip | No overlap | | | Overlap | | |
|---|---|---|---|---|---|---|
| IED[mm]\N ch | 4 | 6 | 8 | 4 | 6 | 8 |
| 5 | 1E4 | 1E6 | 1E7 | 1E4 | 1E6 | 1E7 |
| 10 | 1E8 | 1E7 | 1E7 | 1E6 | 1E7 | 1E7 |

N ch = number of electrodes; IED = Inter-electrode distance

Supplementary Table 4. $k_{Corr\ Opt}$ values for Depths

| Depth | | | | | |
|---|---|---|---|---|---|
| IED [mm]\N ch | 4 | 8 | 10 | 15 | 18 |
| 3 | 1E3 | 1E2 | 1E2 | 1E2 | 1E8 |
| 5 | 1E2 | 1E2 | 1E2 | 1E2 | 1E9 |
| 10 | 1E2 | 1E3 | 1E3 | 1E3 | 1E10 |

N ch = number of electrodes; IED = Inter-electrode distance

In the third round, we evaluated the accuracy of our method given optimal or near-optimal parameters. All constraints were used in the two energy-minimization steps, i.e., the procedure was fully-functional. Between 20 and 30 simulations per condition were evaluated for the optimal ($k_{CorrOpt}$), and near-optimal parameters (0.1 $k_{Corr\ Opt}$ and 10 $k_{Corr\ Opt}$). The latter was to verify the stability of the results. For simplicity, we only report the localization performance using optimal parameters in the final round. Figure 4 shows the accuracy of the localized simulations, and Figure S1 shows the normalized maximum localization error (Eq. 11).

To evaluate the effect of IED, Noise, array Type, and the presence of Overlaps on the normalized median localization error $d_{Loc\ Med}$ (Eq. 10) we performed a multiple linear regression using the following model

$$d_{Loc\ Med} \sim 1 + IED + Noise\ Level + Type + Overlaps + IED*Overlap + Noise*Type + Overlap*Type$$

Similarly, we evaluated the effect of these variables on the normalized maximum localization $d_{Loc\ Max}$ (Eq. 11), i.e.,

$$d_{Loc\ Max} \sim 1 + IED + Noise\ Level + Type + Overlaps + IED*Overlap + Noise*Type + Overlap*Type$$

Supp. Tables 5 and 6 show the N-way ANOVA results for these regression models.

Supplementary Table 5. ANOVA table of effects on mean localization error ($d_{Loc\ Med}$)

| Source | SumSq | DF | MeanSq | F | pValue | Omega$^2$ |
|---|---|---|---|---|---|---|
| IED | 0.573 | 1 | 0.573 | 2827.79 | 0.00E+00 | ***0.235 |
| Noise | 0.234 | 1 | 0.234 | 1157.38 | 1.64E-231 | **0.096 |
| Type | 0.178 | 1 | 0.178 | 880.56 | 2.61E-180 | **0.073 |
| Overlap | 0.072 | 1 | 0.072 | 354.00 | 1.04E-76 | *0.029 |
| Overlap*Type | 0.117 | 1 | 0.117 | 577.57 | 9.31E-122 | **0.048 |
| Noise*Type | 0.045 | 1 | 0.045 | 222.12 | 2.52E-49 | *0.018 |
| IED*Overlap | 0.041 | 1 | 0.041 | 201.59 | 5.27E-45 | *0.017 |
| Error | 1.177 | 5813 | 0.000 | | | |

Asterisks in the Omega$^2$ columns indicate large (0.14, ***), medium (0.06, **), and small (0.01, *) effect sizes (Field, 2013).

Supplementary Table 6. ANOVA table of effects on maximum localization error ($d_{Loc\ Max}$)

| Source | SumSq | DF | MeanSq | F | pValue | Omega$^2$ |
|---|---|---|---|---|---|---|
| Overlap | 3.532 | 1 | 3.532 | 373.606 | 1.00E-80 | **0.058 |
| IED | 0.892 | 1 | 0.892 | 94.341 | 3.91E-22 | *0.015 |
| Noise | 0.489 | 1 | 0.489 | 51.696 | 7.30E-13 | *0.008 |
| Type | 0.221 | 1 | 0.221 | 23.401 | 1.35E-06 | 0.003 |
| Noise*Type | 0.168 | 1 | 0.168 | 17.736 | 2.58E-05 | 0.003 |
| Overlap*Type | 0.196 | 1 | 0.196 | 20.725 | 5.41E-06 | 0.003 |
| IED*Overlap | 0.083 | 1 | 0.083 | 8.729 | 3.14E-03 | 0.001 |
| Error | 54.958 | 5,813 | 0.009 | | | |

Asterisks in the Omega$^2$ columns indicate large (0.14, ***), medium (0.06, **), and small (0.01, *) effect sizes (Field, 2013).

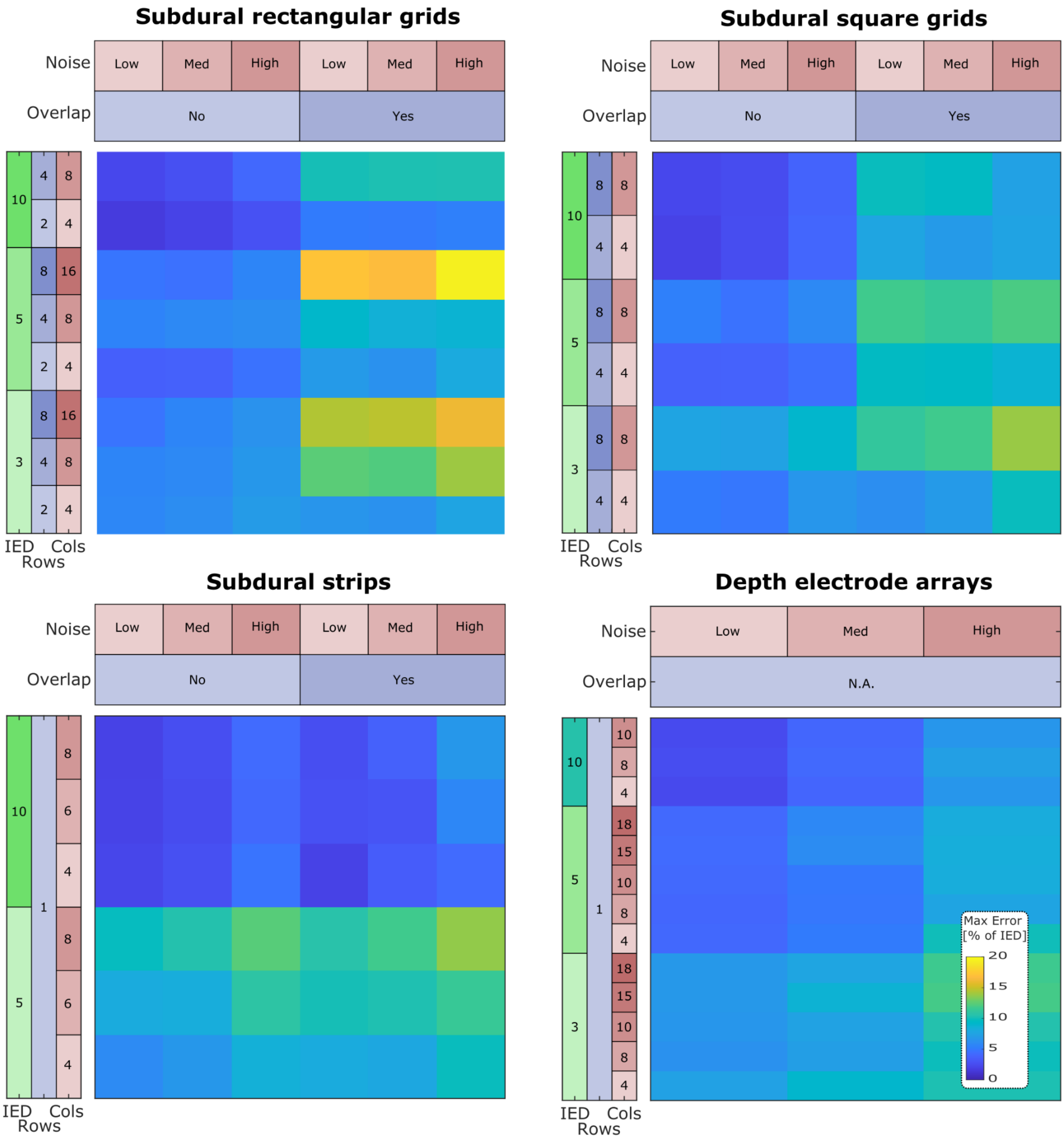

*Figure S1. GridFit maximum localization error of simulated CT artifacts*

### 3. *CEPA* brain-shift compensation supplementary material

#### 3.1. Projection error distance

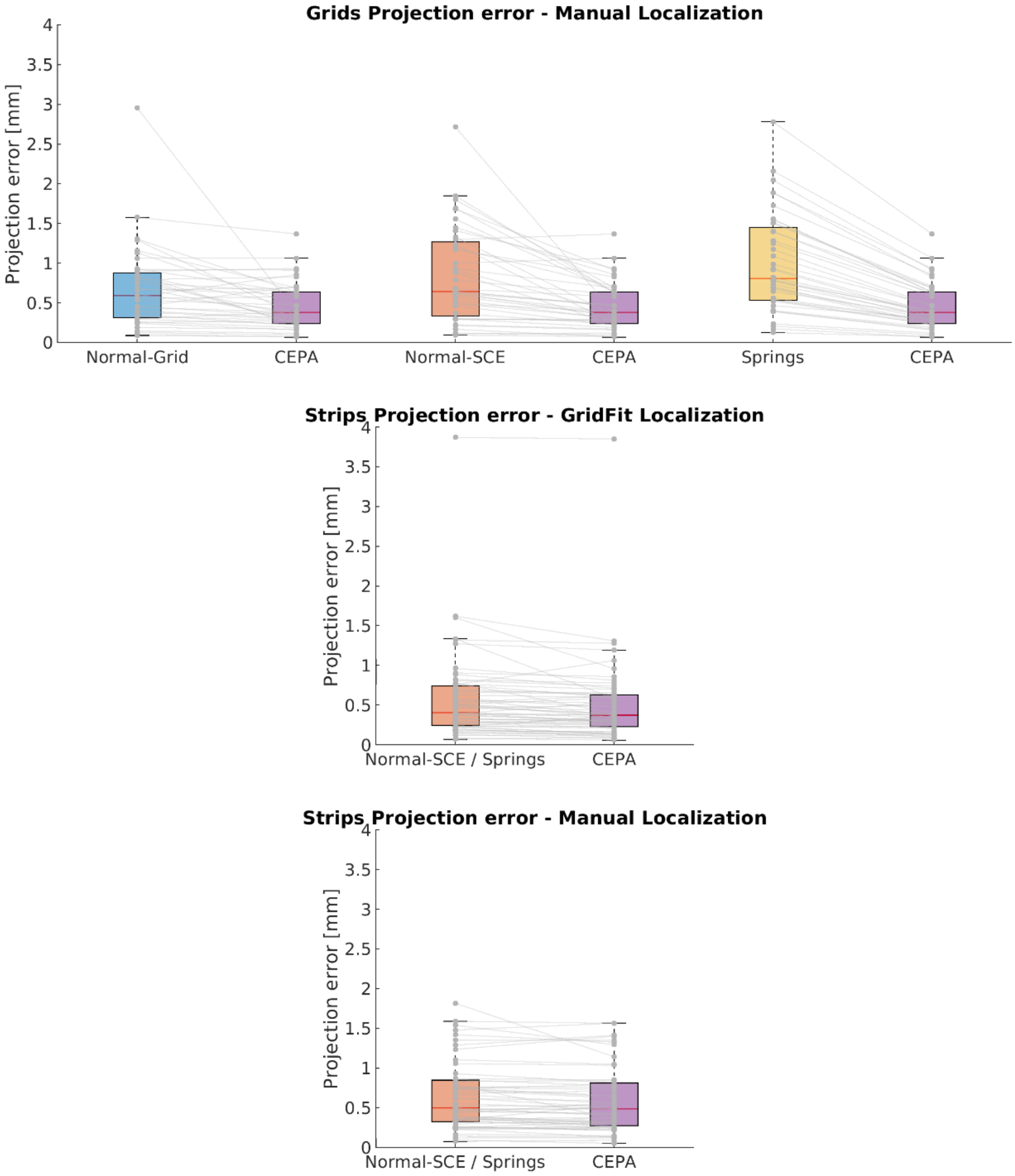

***Figure S2. Projection error comparison of Brain-shift compensation methods***

*Comparison of projection error of Normal-Grid, Normal-SCE, and Springs methods with CEPA.*
*For strip electrodes, the reference location was obtained as the mean between the Normal-SCE and Springs method results. Therefore, the errors for these methods are the same. The normal-Grid method is not available for strips.*

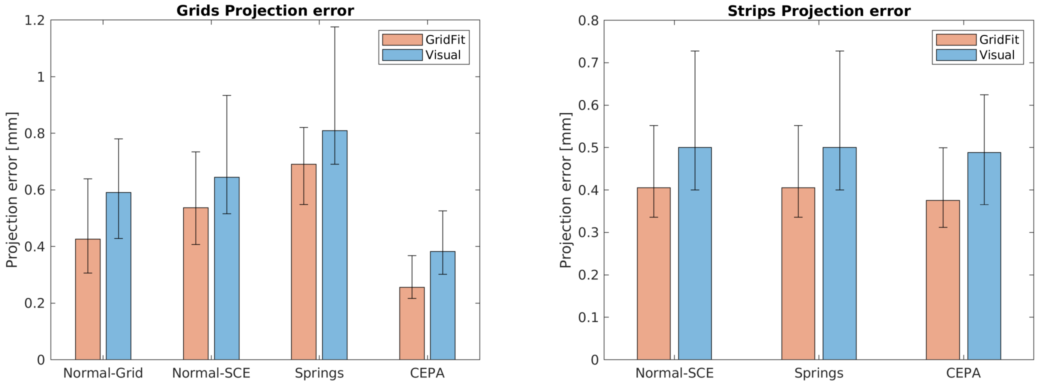

***Figure S3. Projection error comparison of GridFit and visually localized CT artifacts.***
*Comparison of projection error for GridFit and visually localized CT artifacts.*
*It can be observed that visual localization introduces errors that propagate to the back-projected coordinates. Median projection errors were obtained from 40 grids and 59 strips. Error bars denote 95% CI of the median obtained by bootstrapping. Note that statistical tests (Supp. Tables 6 and 7) were done on paired samples and error bars in the plots do not depict errors associated with those tests.*

Supplementary Table 7. *Projection error statistics for electrode grids.*

| *Grids* | | Median projection error [mm] | | | | Comparison CEPA vs. | | | | | |
|---|---|---|---|---|---|---|---|---|---|---|---|
| | | | | | | Normal-Grid | | Normal-SCE | | Springs | |
| | | Normal-Grid | Normal-SCE | Springs | CEPA | z-value | p-value | z-value | p-value | z-value | p-value |
| Loc. | GridFit | 0.43 | 0.54 | 0.69 | 0.26 | 4.01 | 3.01 E-5 | 5.37 | 3.94 E-8 | 5.50 | 1.85 E-8 |
| | Visual | 0.59 | 0.64 | 0.81 | 0.38 | 3.92 | 4.46 E-5 | 5.22 | 8.85 E-8 | 5.50 | 1.85 E-8 |
| GridFit vs. Visual | z-value | 4.57 | 3.09 | 3.77 | 4.67 | | | | | | |
| | p-value | 2.36 E-6 | 9.73 E-4 | 8.15 E-5 | 1.50 E-6 | | | | | | |

Projection error statistics for grids and their comparison with CEPA. Z-values were obtained from the Wilcoxon Signed Rank Test. Results indicate that *CEPA* significantly outperformed the alternative projection methods and that *GridFit* outperformed Visual localization of CT artifacts. Note: Z- and p-values comparing *Spring* vs. *CEPA* from *GridFit* and Visual data are correct and the same by coincidence.

Supplementary Table 8. *Projection error statistics for electrode strips.*

| *Strips* | | Median projection error [mm] | | Comparison CEPA vs. | |
|---|---|---|---|---|---|
| | | | | Normal-SCE / Springs | |
| | | Normal-SCE / Springs | CEPA | z-value | p-value |
| Localization | GridFit | 0.4 | 0.37 | 3.11 | 9.24 E-4 |
| | Visual | 0.5 | 0.49 | 4.502 | 3.36 E-6 |
| GridFit vs. | z-value | 3.40 | 3.60 | | |

| | | | |
|---|---|---|---|
| Visual | p-value | 3.27 E-4 | 1.57 E-4 |

Projection error statistics for strips and their comparison with *CEPA*. Z-values were obtained from the Wilcoxon Signed Rank Test. Results indicate that *CEPA* significantly outperformed the alternative methods. Moreover, the projection of *GridFit* localized CT artifacts outperformed the Visual one.

Note: *Normal-Grid* method is not available for strips. Therefore, the reference location was obtained as the mean between the *Normal-SCE* and *Springs* method results, and the errors and statistical results for these methods were the same.

##### 3.2. Projection roughness

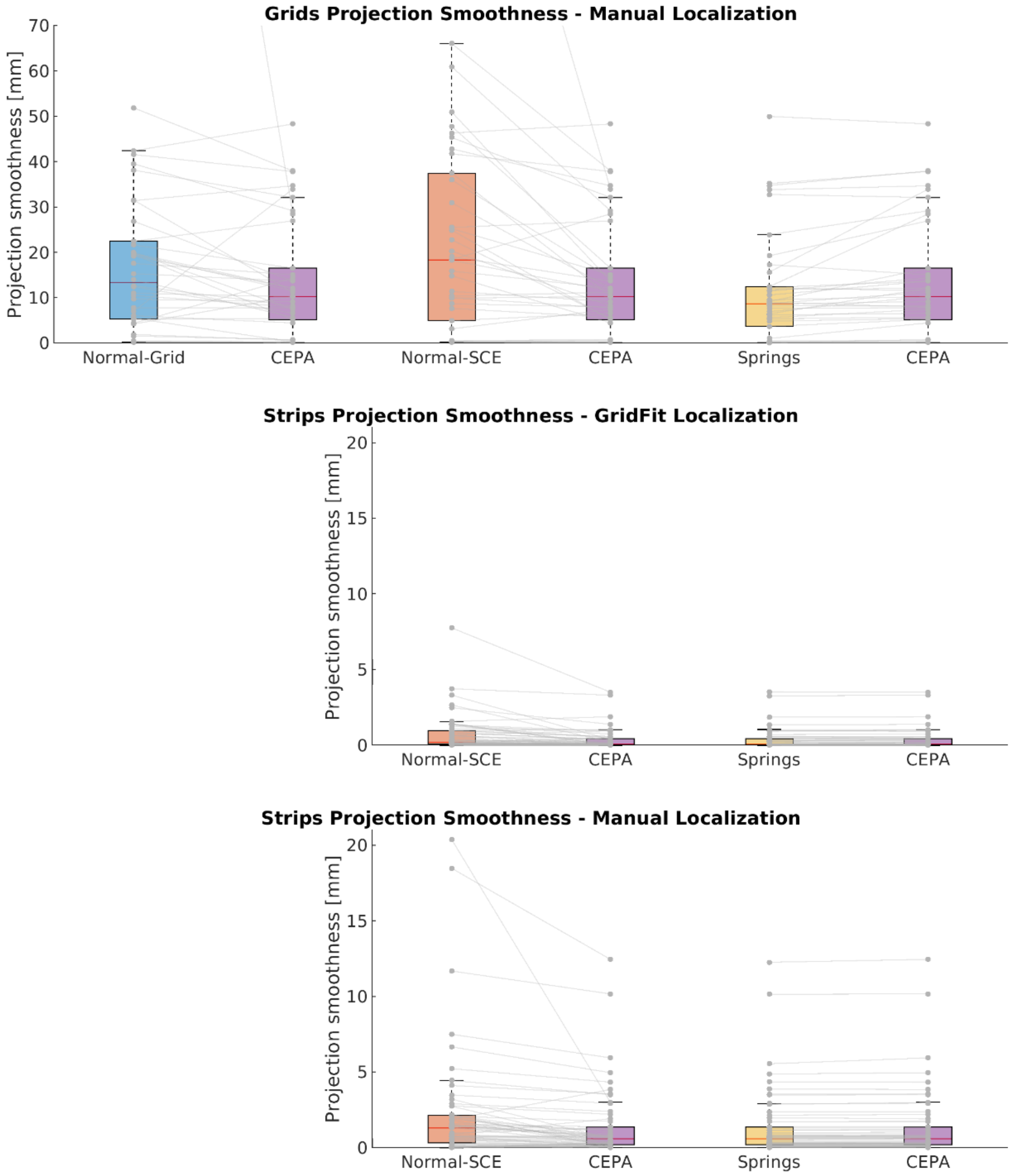

***Figure S4. Roughness comparison of Brain-shift compensation methods.***
*Comparison of projection roughness of Normal-Grid, Normal-SCE, and Springs methods with CEPA.*

All comparisons produced significant statistical differences. *Data were obtained from 40 grids and 59 strips.*

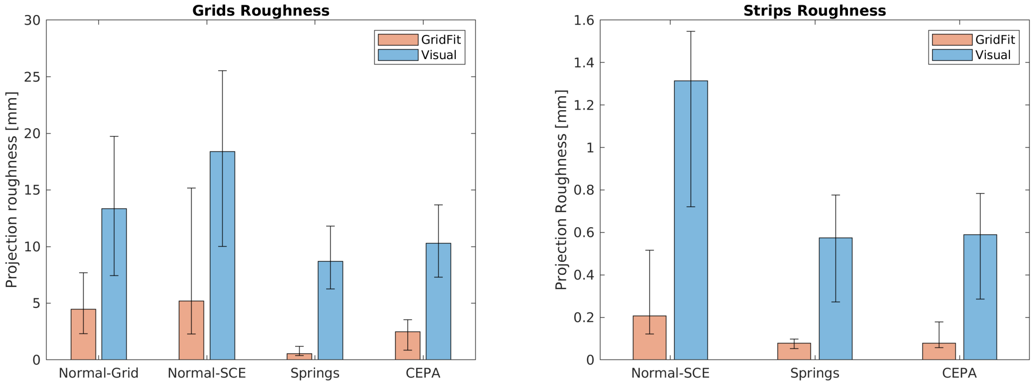

***Figure S5. Roughness comparison of GridFit and visually localized CT artifacts.***
*Comparison of projection roughness for GridFit and visually localized CT artifacts.*
*It can be inferred from the above that visual localization variance in the inter-electrode distance propagates to the back-projected coordinates as reduced roughness.*
*Median projection errors were obtained from 40 grids and 59 strips. Error bars denote 95% CI of the median obtained by bootstrapping.*
*Note that statistical tests (Supp. Tables 9 and 10) were done on paired samples and error bars in the plots do not depict errors associated with those tests.*

Supplementary Table 9. *Projection roughness statistics for grids.*

| *Grids* | | Median roughness [mm] | | | | Comparison CEPA vs. | | | | | |
|---|---|---|---|---|---|---|---|---|---|---|---|
| | | | | | | Normal-Grid | | Normal-SCE | | Springs | |
| | | Normal-Grid | Normal-SCE | Springs | CEPA | z-value | p-value | z-value | p-value | z-value | p-value |
| Loc. | GridFit | 4.47 | 5.21 | 0.55 | 2.48 | 3.90 | 4.79 E-5 | 4.84 | 6.37 E-7 | -4.95 | 3.80 E-7 |
| | Visual | 13.35 | 18.38 | 8.71 | 10.29 | 3.12 | 9.10 E-4 | 3.58 | 1.70 E-4 | -3.83 | 6.44 E-5 |
| GridFit vs. Visual | z-value | 4.31 | 4.73 | 5.29 | 5.21 | | | | | | |
| | p-value | 8.27 E-6 | 1.14 E-6 | 6.01 E-8 | 9.63 E-8 | | | | | | |

*Roughness of projection errors for the different brain-shift compensation methods, and comparison with CEPA. Z-values were obtained from the Wilcoxon Signed Rank Test.*

Supplementary Table 10. *Projection roughness statistics for strips.*

| *Strips* | | Median roughness [mm] | | | Comparison CEPA vs. | | | |
|---|---|---|---|---|---|---|---|---|
| | | | | | Normal-SCE | | Springs | |
| | | Normal-SCE | Springs | CEPA | z-value | p-value | z-value | p-value |
| Loc. | GridFit | 0.21 | 0.08 | 0.08 | 4.01 | 3.01 E-5 | -1.89 | 2.93 E-2 |
| | Visual | 1.31 | 0.57 | 0.59 | 4.52 | 3.02 E-6 | -3.45 | 2.77 E-4 |
| GridFit vs. Visual | z-value | 4.62 | 4.89 | 4.93 | | | | |
| | p-value | 1.96 E-6 | 4.92 E-7 | 4.06 E-7 | | | | |

*Statistical results for roughness comparison of the different methods and comparison with CEPA. Z-values were obtained from the Wilcoxon Signed Rank Test. Normal-Grid is not available for strips.*

##### 3.3. Projection distance vs. local deformation

We studied the local deformations of grids and strips in relationship to the projection distance. More precisely, local deformations were defined as the relationship between the median distance of an electrode $e_j$ with its neighbors $e_k$ normalized by the IED, i. e., median($\{||e_j - e_k||\}$) where k index neighbor electrodes. The projection distance was computed from the localized CT artifact $e_{0j}$ to the back-projected coordinate $e_j$, i.e., $||e_j - e_{0j}||$, normalized by the distance to the brain center.
The effect of the distance was evaluated with linear mixed-effects models (LME) of the deformations, with fixed effects for distance and random effects for distance (and intercept in *Springs* and *CEPA*) grouped by array and patient. Specifically:

$$Deformation \sim Distance + (Distance \mid Patient{:}Array)$$

for *Normal-Grid* and *Normal-SCE* projection methods, and

$$Deformation \sim 1 + Distance + (1 \mid Patient{:}Array) + (Distance \mid Patient{:}Array)$$

for *Springs* and *CEPA* projection methods. Model parameters were estimated using the Maximum Likelihood Estimation method. Including an intercept factor in the last two LME models (*Springs* and *CEPA*) considers that local deformations could occur even when the projection distance is zero. This is because each projected coordinate can be locally deformed by the complete set of projections. In contrast, the first two methods (*Normal-Grid* and *Normal-SCE*) should not produce this effect.
Data outliers from each regression were detected using Grubbs's test implemented in Matlab's function "isoutlier". Outliers from all models (*Normal-Grid* 49, *Normal-SCE* 36, *Springs* 38, and *CEPA* 19) were combined (N = 114) and removed to perform the final data regressions (out of N = 2327).
Fewer samples were available for the *Normal-Grid* method since it is not applicable for strips. Only data obtained from localizing CT artifacts with the *GridFit* algorithm were analyzed for simplicity.

T-tests were performed to determine the significance of fixed effects, and Likelihood Ratio Tests (LRT) were used to compare the LME models. LRT showed negative results for including correlated random effects in the second model.
LRT showed positive results when models were compared against simpler linear regressions with no random effects. Along the same line, the confidence intervals of the random effects excluded zero.
It can be observed in Figure 6E that all methods except *Springs* were sensitive to the projection distance. *Normal-Grid* showed the strongest effect of distance over deformation, followed by *CEPA* and *Normal-SCE*. Moreover, variable levels of dispersion were observed for the different projections as indicated by the adjusted $R^2$.
Supplementary Table 11 shows modeling details.

Supplementary Table 11. S*ummary of the linear mixed-effects modeling of Deformation*

| *Method* | $\beta_0$ | $\beta_1$ | N obs | SE | t-value fixed effect | p-value fixed effect | Adj. $R^2$ |
|---|---|---|---|---|---|---|---|
| Normal-Grid | N.A. | 0.31 | 1960 | 0.072 | 4.33 | 1.53 E-5 | 0.30 |
| Normal-SCE | N.A. | 0.67 | 2213 | 0.092 | 7.44 | 1.46 E-13 | 0.72 |
| Springs | -1.58 | -0.03 | 2213 | 0.032 | 1.02 | 0.31 | 0.58 |
| CEPA | 0.39 | 0.36 | 2213 | 0.068 | 5.34 | 9.93 E-8 | 0.74 |

*Summary of the LME modeling of the local deformation as a function of the projection distance for the different projection methods. DF: Degrees of Freedom; SE: Standard Error*

##### 3.4. Alternative reference location

In the main article, we described the localization errors in relation to a mean location reference, i.e., the mean of *Normal-SCE*, *Normal-Grid*, and *Springs* methods' results. Even if this is a parsimonious approach, it might not reflect the best possible back-projection. Here, we explored the possibility that any of the evaluated methods could be more reliable as a reference.
Therefore, each method was defined as a reference, and the errors were computed against these locations. The figure below (Fig S7) shows the median errors of using each method as a reference location. Observe that in all situations but one (*Springs-ref*, *CEPA* vs. *Normal-Grid*), the *CEPA* method statistically outperformed the other evaluated methods. These results suggest that in the worst scenario assessed here, the *CEPA* method produced the second-best localization alternative, with a median error below 1.3mm in all cases. See statistical details in Supplementary Tables 12 and 13.

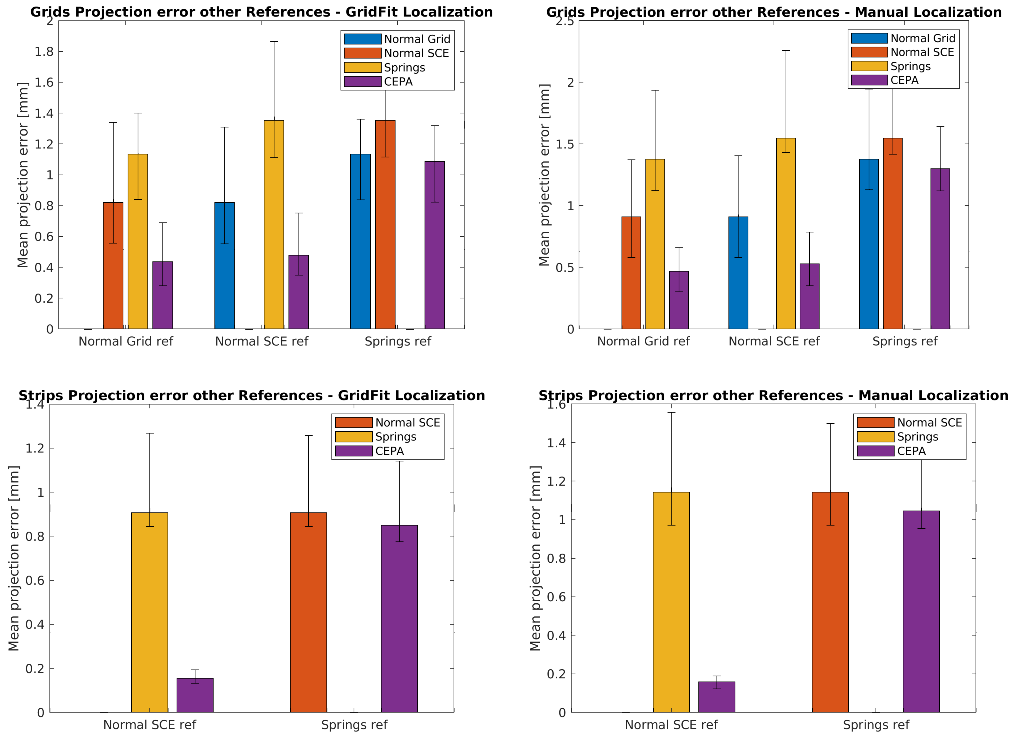

***Figure S7. Projection error when selecting other brain-shift compensation methods as a reference.***
*In all situations but one (Springs-ref, CEPA vs. Normal-Grid), the CEPA method statistically outperformed the other methods evaluated.*
*Wilcoxon signed-rank test was used to assess pairwise statistical significance between methods. However, for illustration simplicity, only medians and CI are shown.*
*Median projection errors were obtained from 40 grids and 59 strips. Error bars denote 95% CI of the median obtained by bootstrapping.*
*Note that statistical tests (Supp. Tables 12 and 13) were done on paired samples and error bars in the plots do not depict errors associated with those tests.*

Supplementary Table 12. *Projection error statistics for grids using alternative references.*

| *Grids* | | Normal-Grid Ref | | | Normal-SCE Ref | | | Springs Ref | | |
|---|---|---|---|---|---|---|---|---|---|---|
| | | CEPA | Normal-SCE | Springs | CEPA | Normal-Grid | Springs | CEPA | Normal-Grid | Normal-SCE |
| GridFit | error [mm] | 0.44 | 0.82 | 1.13 | 0.48 | 0.82 | 1.35 | 1.09 | 1.13 | 1.35 |
| | z-value | | 5.50 | 5.29 | | 5.50 | 5.38 | | 0.71 | 5.50 |
| | p-value | | 1.85 E-8 | 6.14 E-8 | | 1.85 E-8 | 3.66 E-8 | | 0.24 | 1.85 E-8 |
| Visual | error [mm] | 0.47 | 0.91 | 1.38 | 0.53 | 0.91 | 1.55 | 1.30 | 1.38 | 1.55 |
| | z-value | | 5.50 | 5.49 | | 5.50 | 5.44 | | 0.25 | 5.18 |
| | p-value | | 1.85 E-8 | 2.00 E-8 | | 1.85 E-8 | 2.71 E-8 | | 0.40 | 1.10 E-7 |

*Localization errors of CEPA vs. Normal-Grid, Normal-SCE, and Springs, when one of these alternative methods was used as reference. Statistical results were obtained from the Wilcoxon signed-rank test, contrasting CEPA with other methods. The error row refers to the median projection error.*

Supplementary Table 13. *Projection error statistics for strips using alternative references.*

| *Strips* | | Normal-SCE Ref | | Springs Ref | |
|---|---|---|---|---|---|
| | | CEPA | Springs | CEPA | Normal-SCE |
| GridFit | error [mm] | 0.16 | 0.91 | 0.85 | 0.91 |
| | z-value | | 6.68 | | 6.67 |
| | p-value | | 1.23 E-11 | | 1.29 E-11 |
| Visual | error [mm] | 0.16 | 1.14 | 1.05 | 1.14 |
| | z-value | | 6.68 | | 6.38 |
| | p-value | | 1.23 E-11 | | 8.75 E-11 |

*Localization errors of CEPA vs. Normal-Grid, Normal-SCE, and Springs, when one of these alternative methods was used as reference. Statistical results were obtained from the Wilcoxon signed-rank test, contrasting CEPA with the other methods. The error row refers to the median projection error.*

##### 3.5. Linear Mixed-Effect modeling of HFA activity by attenuation models

To assess the quality of the different back-projection algorithms, we compared the explanatory power of their respective *Estimated HFA* patterns predicting resting-state *Measured HFA*. We used the following LME model:

$$measured\_HFA \sim 1 + estimated\_HFA + (1 \mid Patient{:}Array)$$

where *measured_HFA* is the mean HFA obtained from the intracranial recordings, estimated*_HFA* is the expected HFA attenuation given the projection method (i.e., $estimated\_HFA_{Normal\text{-}Grid}$, $estimated\_HFA_{Normal\text{-}SCE}$, $estimated\_HFA_{Springs}$, or $estimated\_HFA_{CEPA}$), *Array* is the array number, and *Patient* is the patient number. *estimated_HFA* patterns were computed as the negative log-transformed distance between the projected electrodes and the pial surface. Log transformation was applied to correct left-skew data.

Data were modeled using projections from *GridFit* and visually localized CT artifacts. Parameters were estimated using the Maximum Likelihood Estimation method. Noisy or epileptic channels were removed. Therefore, the total number of observations for all models was 487 (out of the initial 528 channels). Supplementary Tables 14 and 15 below show modeling details for each case.

T-tests were performed to determine the significance of fixed effects, and Likelihood Ratio Tests (LRT) were used to compare the LME models. LRT supported the actual model against simpler models with no random effects. Along the same line, the confidence intervals of the random effects excluded zero.

Bayesian Information Criterion was computed given likelihood values *L* obtained by fitting competing models to data, the corresponding number of estimated model parameters *numParam*, and the corresponding sample sizes used in estimation *numObs*:

$$BIC = -2*logL + log(numObs)*numParam$$

Supplementary Table 14. *Summary of the linear mixed-effects modeling of HFA using GridFit localized coordinates.*

| Model | β | SE | t-value fixed effect | p-value fixed effect | Adj. $R^2$ | BIC |
|---|---|---|---|---|---|---|
| Normal-Grid | 2.19E-01 | 3.22E-02 | 6.79 | 3.29E-11 | 0.475 | 39.82 |
| Normal-SCE | 2.16E-01 | 3.16E-02 | 6.82 | 2.75E-11 | 0.476 | 39.61 |
| Springs | 2.05E-01 | 3.34E-02 | 6.15 | 1.65E-09 | 0.467 | 47.40 |
| CEPA | 2.15E-01 | 3.17E-02 | 6.79 | 3.41E-11 | 0.476 | 39.95 |

*SE: Standard Error, BIC: Bayesian Information Criterion.*

Supplementary Table 15. *Summary of the linear mixed-effects modeling of HFA using visually localized coordinates.*

| Model | β | SE | t-value fixed effect | p-value fixed effect | Adj. $R^2$ | BIC |
|---|---|---|---|---|---|---|
| Normal-Grid | 2.18E-01 | 3.29E-02 | 6.63 | 8.87E-11 | 0.473 | 41.74 |
| Normal-SCE | 2.18E-01 | 3.22E-02 | 6.78 | 3.52E-11 | 0.476 | 40.13 |
| Springs | 1.56E-01 | 3.28E-02 | 4.76 | 2.58E-06 | 0.449 | 61.33 |
| CEPA | 2.15E-01 | 3.24E-02 | 6.63 | 9.19E-11 | 0.473 | 41.90 |

*SE: Standard Error, BIC: Bayesian Information Criterion.*
